# Functional coupling of the mesencephalic locomotor region and V2a reticulospinal neurons driving forward locomotion

**DOI:** 10.1101/2022.04.01.486703

**Authors:** Martin Carbo-Tano, Mathilde Lapoix, Xinyu Jia, François Auclair, Réjean Dubuc, Claire Wyart

**Affiliations:** Sorbonne Université, Institut du Cerveau (Paris Brain Institute, ICM), Inserm U 1127, CNRS UMR 7225, APHP, Hôpital Pitié-Salpêtrière, Paris, France; Université de Paris, Paris, France; Department of Neuroscience, Université de Montréal, Montréal, Québec, Canada; Groupe de Recherche en Activité Physique Adaptée, Department of Exercise Science, Université du Québec à Montréal, Montréal, Québec, Canada

**Keywords:** Mesencephalic locomotor region (MLR), locomotion, reticulospinal neurons (RSNs), V2a neurons, *vsx2* transcription factor, brainstem, zebrafish

## Abstract

Locomotion in vertebrates relies on high brain centers converging onto the mesencephalic locomotor region (MLR). How the MLR recruits brainstem reticulospinal neurons (RSNs) to initiate locomotion is incompletely understood due to the challenge of recording these cells in vivo. To tackle this question, we leveraged the transparency and genetic accessibility of larval zebrafish. In this model organism, we uncovered the locus of the MLR as a small region dorsal to the locus coeruleus containing glutamatergic and cholinergic neurons. MLR stimulations reliably elicited forward bouts of controlled duration and speed. We find that the MLR elicits forward locomotion by recruiting V2a RSNs in the pontine and retropontine regions, and gradually in the medulla. Remarkably, recruited V2a RSNs in the medulla act as maintain cells encoding speed of forward locomotion. Altogether, our study reveals that the MLR recruits genetically-identified reticulospinal neurons in the medulla to control the kinematics of exploration.

## Introduction

Locomotion is the fundamental behavior enabling animals to move in their environment, and thereby to rapidly adapt to the challenges of their environment while satisfying their inner needs. Locomotion can be sensory-evoked to avoid a threatening stimulus for example during escape responses, or goal-directed to explore the environment in search for resources. Motor circuits in vertebrate species are conserved at the anatomical, physiological, and molecular levels^1–3^. The production of locomotor movements relies on the recruitment of command neurons in the brainstem, referred to as reticulospinal neurons (RSNs) in the reticular formation. RSNs integrate synaptic inputs from higher brain areas and from the periphery, and in turn, instruct the spinal circuits that produce movements^4,5^ (for reviews, see^6–8^). The RSNs have been shown to play a critical role in starting, maintaining and stopping locomotion^5,9–11^, as well as in adjusting posture and steering^12–14^.

A critical region involved in the production of locomotion upstream the reticular formation is the mesencephalic locomotor region (MLR), first identified in decorticated cats^15^. The MLR has since been functionally defined in numerous other vertebrate species including lampreys^16^, salamanders^17^, rats^18^, mice^19^, rabbits^20^, guinea pigs^21^, pigs^22^ and monkeys^23^. Electrical stimulation of the MLR triggers forward locomotion in a graded fashion as a function of the stimulation intensity^15,20^ (for review, see^24^). The conservation of MLR properties across vertebrate species suggests that this brainstem structure is essential to vertebrate locomotion.

Anatomical studies indicate that MLR is localized in the vicinity of the mesopontine cholinergic nuclei (for review, see^24^) and corresponds in mammals to the pedunculopontine nucleus^25^ (PPN) and the cuneiform nucleus^15^ (CuN). In all vertebrate species investigated, the MLR induces locomotion via the activation of RSNs^26–33^. However, the MLR-dependent recruitment pattern of RSNs that elicits forward locomotion has not yet been resolved due to the complexity and difficulty to access the entire reticular formation *in vivo*. An interesting subclass of RSNs downstream of the MLR expresses the transcription factor *vsx2+* and corresponds to V2a neurons^34–36^. While V2a RSNs have been shown to be necessary for the initiation of locomotion^37^, some studies have highlighted the role of pontine and retropontine V2a RSNs in turning^38–40^ and stopping locomotion^41^. Yet, the nature of V2a RSNs recruited by the MLR to trigger forward movement at various locomotor speeds is unknown.

Here we leverage the transparency and genetic accessibility of larval zebrafish to unravel how genetically identified RSNs are recruited by the MLR to produce forward locomotion. We first identified functionally and anatomically the location of the MLR in larval zebrafish. We then combined behavioral recordings and population calcium imaging to investigate the recruitment of V2a RSNs across the brainstem upon MLR stimulation. We found that subsets of brainstem V2a RSNs are gradually recruited upon MLR stimulation at increasing intensities. Our study uncovers that a population of V2a reticulospinal neurons in the medulla is sustainably recruited during MLR-evoked forward locomotion and encodes key parameters controlling speed of forward locomotion.

## Results

### Functional identification of the MLR in larval zebrafish

Previous studies in fish found that electrical stimulation of the dorsocaudal tegmentum in the midbrain that contains the nucleus of the medial lateral fasciculus (nMLF) induced locomotion^42–46^, leading some authors to suggest this region corresponds to the MLR^42,43,46,47^. However, the nMLF is adjacent to the oculomotor nuclei (N3) and not to the mesopontine cholinergic nuclei where the MLR has been identified in all other vertebrates^24^ (**Fig. 1a-b**). Based on previous anatomical and physiological experiments performed in other vertebrate species (summarized in **Fig. 1a**), including mice^48^, lampreys^16,49^ and salamanders^17^, we localized the homologous mesopontine cholinergic nuclei of larval zebrafish in the prepontine region, medial to the locus coeruleus (**Fig. 1b**).

**Fig. 1.**
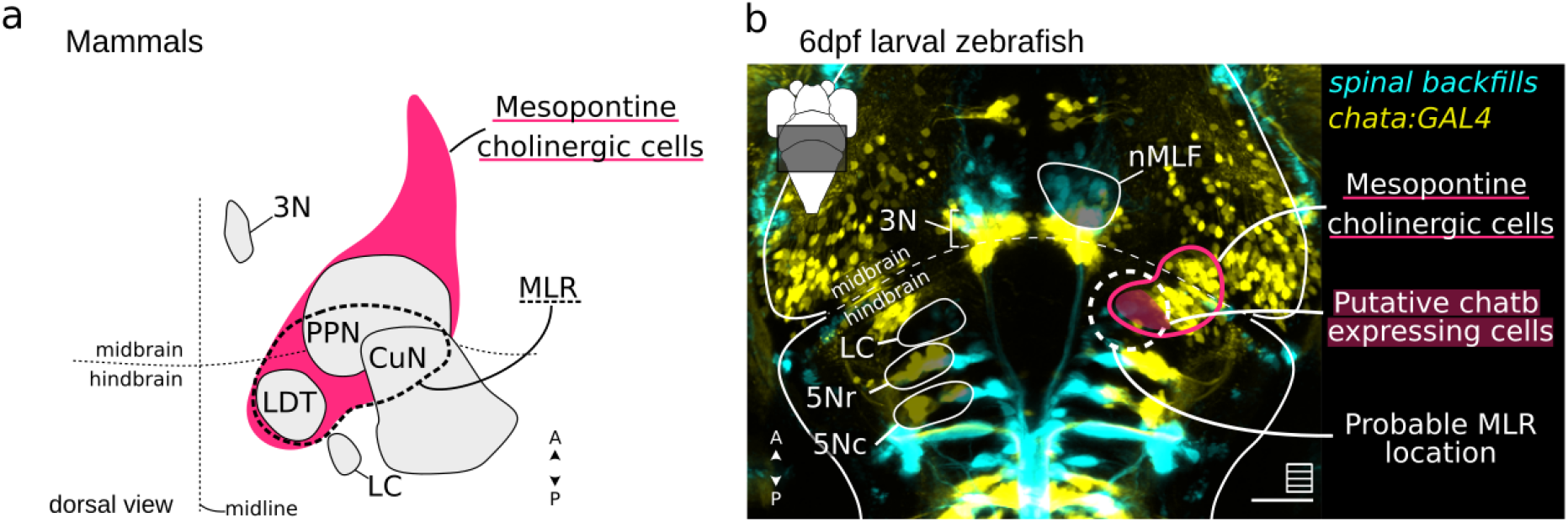
The putative location of the MLR in zebrafish was inferred from previous studies in mammals, amphibians and lampreys showing overlap with mesopontine cholinergic nuclei. **a,** Schematic illustration of the MLR location in mammals (dashed line) in relation to the mesopontine cholinergic cells (magenta). The structures classically defined as part of the MLR are the pedunculopontine nucleus (PPN), cuneiform nucleus (CuN) and laterodorsal tegmental nucleus (LDT). The characteristics of the structures were taken from the Allen Mouse Brain Atlas (accessible at https://atlas.brain-map.org) and the location and extent of the mesopontine cholinergic nucleus was approximated from^48^. **b,** Larval zebrafish brain image depicting the probable MLR location (dashed white line) in relation to the locus coeruleus (LC) and the mesopontine cholinergic cells labeled in the *Tg(chata:GAL4)* transgenic line. The hindbrain reticulospinal neurons and the nMLF are labeled by spinal backfills. All images were taken from the web-interface of *mapzebrain* (https://fishatlas.neuro.mpg.de/, download: 12 Apr. 2020). Sale bar is 50 μm.

In order to narrow down the locus of MLR, we next investigated the behavioral responses elicited by electrical stimulation applied using monopolar tungsten microelectrode inserted in different locations within the mesopontine cholinergic nuclei of larval zebrafish (**Fig. 1b**). Using a high speed camera, we monitored the behavioral responses of 6 days post fertilization (dpf) head-embedded tail-free larvae to 2 to 40 s-long stimulation trains of 2 ms duration pulses occurring at variable frequency (5 - 20 Hz) and intensity (0.1 - 2 μA) (**Fig. 2a,** see **Methods**). The behavioral responses consisted of locomotor bouts corresponding either to coordinated symmetric forward swims (**Fig. 2b**, blue trace), uncoordinated asymmetric tail bends resembling struggle (**Fig. 2b**, magenta trace), or a mix of both occurring sequentially as distinct motor episodes within the same bout (**Fig. 2c**, left). We manually segmented and classified each episode as either forward swim or struggle (see **Methods**). The maximal tail bend amplitude and the median tail beat frequency best differentiated forward swimming from struggle episodes (**Fig. 2c**, right). For each stimulation, we defined a *forward index* as the normalized ratio of forward over struggle elicited episodes (see **Methods**, **Fig. 2d**). For each microelectrode position, we calculated the median *forward index* over all stimulations and then mapped each 3D position to a reference brain space based on the *mapzebrain* atlas^50^ (see **Methods**, **Fig. 2e**). We defined the MLR as the brainstem region comprising all stimulation sites with a positive median forward index. The MLR locus corresponded to a confined area of ~30 μm in radius in rhombomere 1, which was medial and dorsal to the locus coeruleus (LC) (**Fig. 2e, and Supplementary Video 1**). The stimulation of the MLR at 10 Hz was the most effective at inducing forward swimming, while 5 Hz or 20 Hz stimulation frequency often resulted in struggle-like behaviors (**Extended Data Fig. 1**). This observation is consistent with previous results in lampreys and salamanders showing an optimal frequency of stimulation between 2 and 10 Hz^16,17,51^. Altogether, our approach functionally identified the locus of the MLR of larval zebrafish in a prepontine area that is medial to the locus coeruleus in the vicinity of the mesopontine cholinergic nuclei.

**Fig. 2.**
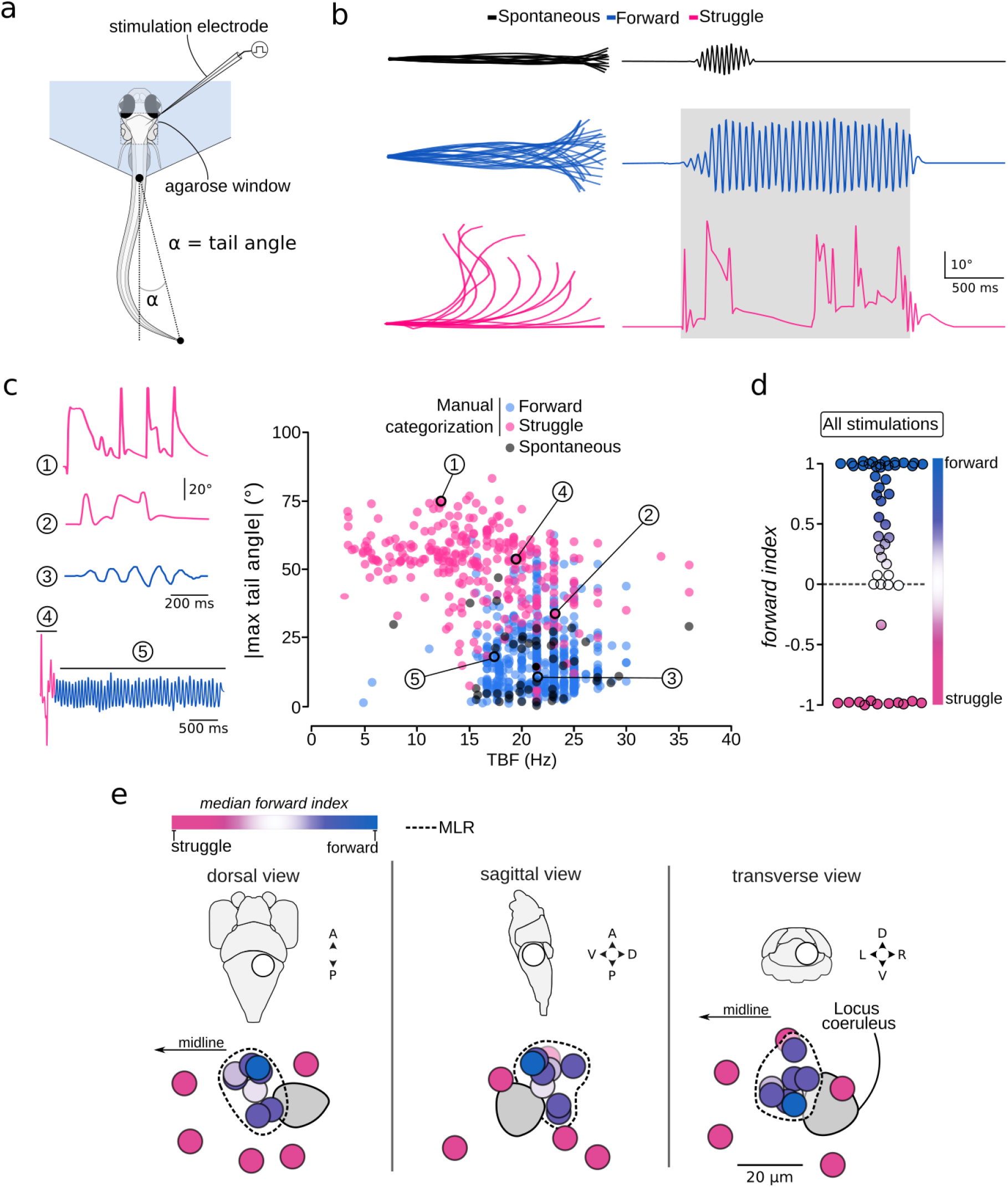
Electrical stimulation of the MLR induces symmetric forward swimming. **a**, Schematic illustration of the behavioral experiments with MLR stimulations **b**, Typical behavioral responses shown as superimposed images of the larval zebrafish tail (left panel), and corresponding tail angle trace (right panel), highlighting differences in symmetry and amplitude between a typical spontaneous swim episode (black, symmetrical and low amplitude tail bends), and electrically-evoked episodes that typically corresponded to a forward swim (blue, symmetrical and moderate amplitude) or to a struggle (magenta, asymmetrical and large amplitude). The gray bar indicates the stimulation duration. **c**, *Left*, representative example of tail angle traces. All locomotor episodes elicited by electrical stimulation were manually segmented and classified as either forward swims (blue) or struggles (magenta) (13 fish, 164 stimulations, 786 episodes). *Right*, values of maximum absolute tail angle versus median tail beat frequency (TBF) for all segmented episodes. Spontaneous episodes are shown as black dots. **d**, Distribution of *forward index* for all simulations applied: *forward index* = (number of forward episodes - number of struggles) / (total number of elicited episodes during stimulation). **e**, Brainstem location of all stimulation sites investigated: color code represents the median *forward index* calculated from all the stimulation applied for each stimulation site (pink, struggle, and blue forward swim, as depicted in panel **d**). The dashed line represents the MLR location covering all stimulation sites with a median forward index above 0. A, anterior; P, posterior; D, dorsal; V, ventral; L, left; R, right. Scale bar is 20 μm.

**Extended Data Fig. 1.**
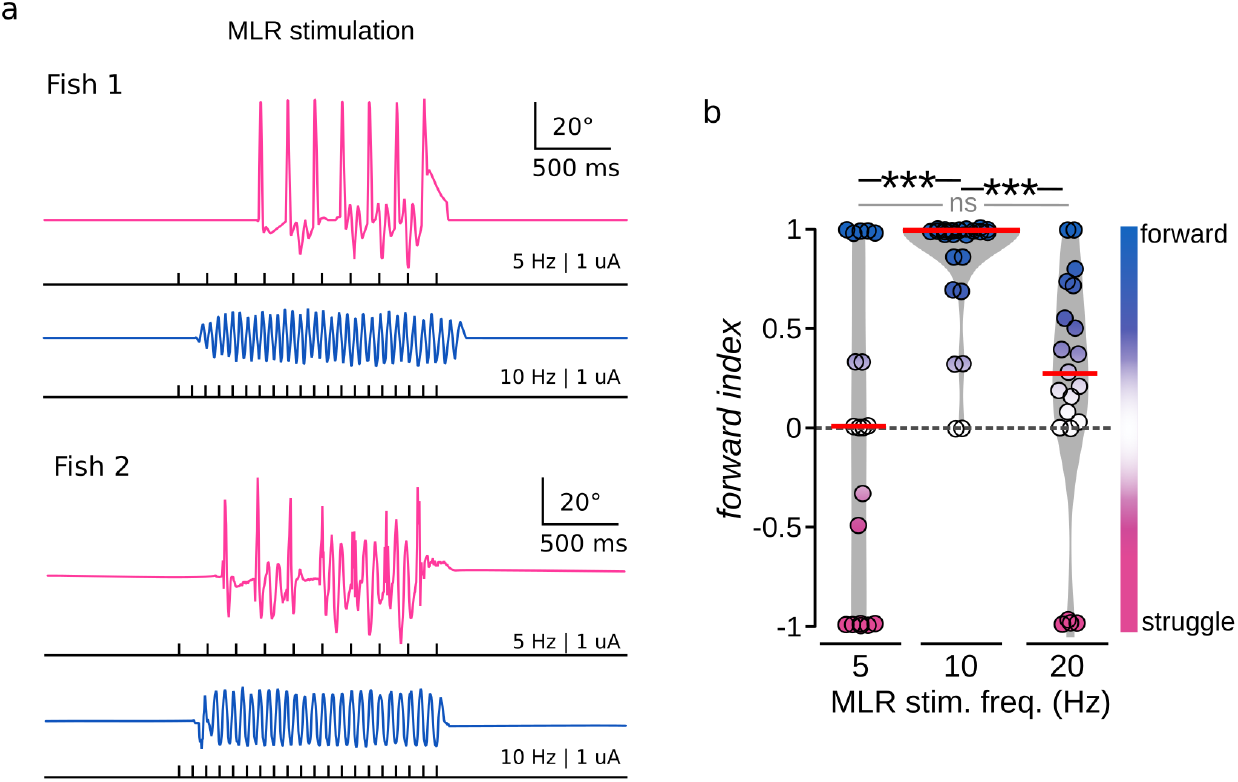
**a,** Representative behavioral responses to MLR stimulation at different frequencies in two different larvae. **b**, *Forward index* for all the stimulation sites according to the stimulation frequency (10 fish and 64 stimulation trials; 5 Hz: N = 19 stimulations, −0.06 ± 0.8, 10 Hz: N = 26 stimulations, 0.919 ± 0.24, 20 Hz: N = 19 stimulations, 0.27 ± 0.55 s; Kruskal-Wallis test, χ^2^ = 105.85, df = 3, p < 0.001, Wilcoxon’s test pairwise comparisons: 5 Hz vs 10 Hz: p < 0.001; 5 Hz vs 20Hz: p = 0.17; 10 Hz vs 20 Hz: p < 0.001).

### Parameters of the MLR stimulation determine the latency, duration, and frequency of locomotion

In previously investigated vertebrate species, the intensity of the MLR stimulation controlled the duration and power of locomotion^16,17,22,27^. We investigated whether the intensity, train duration and pulse frequency defining the MLR stimulation modulated the duration and kinematics parameters of forward swims. In head-fixed tail-free larvae, stimulating the MLR for 2 or 4 s elicited sustained episodes lasting the stimulation train (**Fig. 3a-b,** 10 Hz pulse train at 1 μA; mean ± sd for 2 s long train: 1.91 ± 0.44 s, for 4 s long train: 3.65 ± 0.31 s). In comparison, spontaneous forward swims recorded between stimulation trials only lasted for a few hundreds of milliseconds (**Fig. 3a-b,** spontaneous: 0.41 ± 0.15 s). Furthermore, as previously observed in lampreys and salamanders^16,17^, the delay to locomotor onset was inversely proportional to the stimulation intensity (**Fig. 3c,** 0.1 μA: 1.22 ± 1.97 s; 1 μA: 0.15 ± 0.11 s). We noticed that MLR-induced forward swims exhibited larger median tail bend amplitude (TBA) than spontaneous swims (**Fig. 3d**, spontaneous: median TBA = 5.9 ± 3.16°, MLR-induced: median TBA = 8.5 ± 3.8°), but similar median tail beat frequencies (TBFs) (**Fig. 3e** spontaneous: median TBF = 21.6 ± 3.1 Hz; MLR-induced 21.3 ± 2.9 Hz). Upon closer inspection, we noticed that the TBF probability density distribution upon MLR stimulation was bimodal (**Fig. 3e,** one peak at 17.5 Hz and the other 23.2 Hz, red lines), possibly reflecting different responses to the frequency used for MLR stimulation. We found indeed that MLR-induced forward swims exhibited a median TBF around 17 Hz when the stimulation occurred at frequencies below 15 Hz (**Fig. 3f**, 5 Hz: 17.54 ± 1.31 Hz; 10 Hz: 18.7 ± 2.34 Hz; 15 Hz: 17.66 ± 1.07 Hz) while 20 Hz MLR stimulation elicited forward swims at 23 Hz (**Fig. 3f**, 23.1 ± 1.62 Hz). Altogether, our observations indicate that the properties of the MLR stimulation impacts the duration, the time onset, the amplitude, and the locomotor frequency of forward swims.

**Fig. 3.**
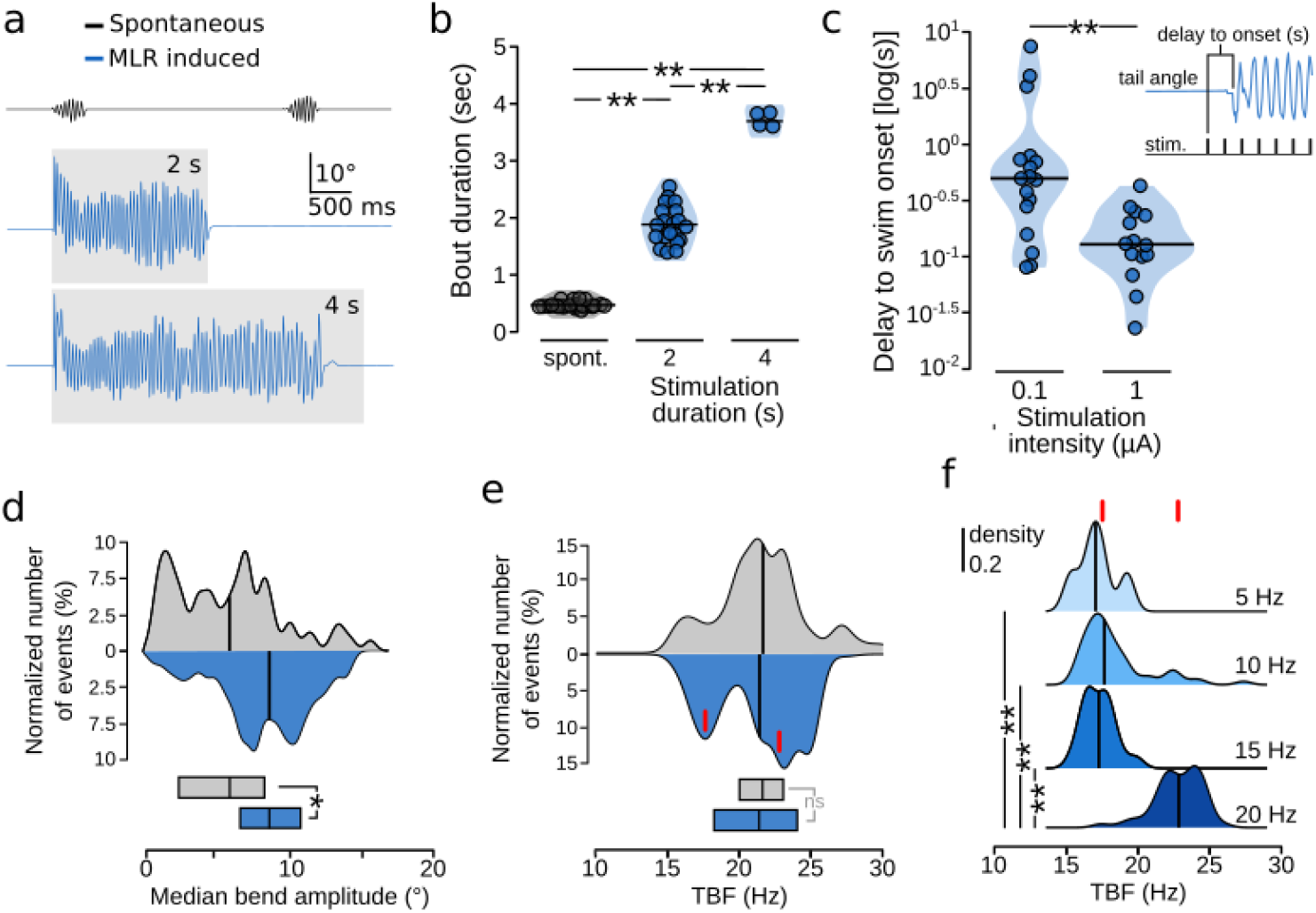
MLR stimulation parameters set the duration, the time onset, the amplitude and the locomotor frequency of forward swims. **a**, Typical example of tail angle traces for spontaneous forward swims (black, top trace) and forward swims induced by stimulating the MLR at 10 Hz, 1 μA for either 2 s (blue, middle trace) or 4 s (blue, bottom trace). The gray box indicates the duration of the train. **b**, Quantification of the duration of forward swims according to the duration of the MLR stimulation (10 Hz, 1 μA), displayed as a violin plot with black line indicating the median value (8 fish; spontaneous: 27 episodes, 0.41 ± 0.15 s, 2 s long stimulation: 19 episodes, 1.91 ± 0.44 s, 4s-long stimulation: 4 episodes, 3.65 ± 0.315 s; Kruskal-Wallis test, x^2^= 38.6, df = 2, p < 0.001, Wilcoxon’s test pairwise comparisons: spont. vs 2 s: p < 0.001; spont. vs 4 s: p < 0.001; 2 s. vs 4 s: p < 0.001). **c**, The delay to swim onset decreased with higher MLR stimulation intensities. Stimulation frequency was set to 10 Hz and duration to 2 s (9 fish; 0.1 μA: 17 episodes, 1.22 ± 1.97 s; 1 μA: 14 episodes, 0.15 ± 0.11 s; Mann-Whitney U Test, W = 199, p < 0.001). **d**, MLR-evoked forward swims reliably exhibited larger tail bend amplitude compared to spontaneous forward swims (12 fish. spont: 67 episodes, 5.9 ± 3.16°; MLR stim: 172 episodes, 8.5 ± 3.8°; Mann-Whitney U Test, W = 8102, p < 0.001). Note the absence of the lowest bend amplitude values when stimulating the MLR. In all panels, the black line indicates the median value. **e**, The distribution of median tail beat frequency (TBF) upon MLR stimulation was bimodal with a first peak at 17.5 Hz and a second at 23.2 Hz (red line represents the peak of the distributions, *normalmixEM* function from R *mixtools* package^52^): median TBF was indistinguishable for forward swims occurring spontaneously or upon MLR stimulation at 5 and 20 Hz (11 fish; spont: 57 episodes, 21.6 ± 3.1 Hz; MLR stim: 242 episodes, 21.3 ± 2.9 Hz; Mann-Whitney U Test, W = 6721, p = 0.7; the boxes at the bottom show the median and 25-75 % quantiles). **f**, 20 Hz MLR stimulation elicited swimming at higher frequency (23 Hz) than during spontaneous or lower frequency of MLR stimulation (~17Hz) (11 fish. 5 Hz: 10 episodes, 17.54 ± 1.31 Hz; 10 Hz: 60 episodes, 18.7 ± 2.34 Hz; 15 Hz: 18 episodes, 17.66 ± 1.07 Hz; 20 Hz: 154 episodes, 23.1 ± 1.618 Hz. Kruskal-Wallis test, χ^2^ = 6237, df = 31, p <0.001, Wilcoxon’s test pairwise comparisons: 5 Hz vs 10 Hz: p = 0.3; 5 Hz vs 15 Hz: p = 0.9; 5Hz vs 20 Hz: p <0.001; 10 Hz vs 15 Hz: p = 0.2; 10 Hz vs 20 Hz: p <0.001; 15 Hz vs 20 Hz: p <0.001). The red lines represent the peak values for each of the distributions from panel (e).

### Neurons in the MLR locus project to the reticular formation

The locomotor output elicited in response to electrical stimulation of the MLR is due to projections onto hindbrain reticulospinal neurons (RSNs)^4,31,32^. To test whether the MLR region contains neurons that project to RSNs in larval zebrafish, we performed retrograde tracing experiments via a unilateral injection of biocytin in the reticular formation between rhombomeres 2 and 6 (**Fig. 4a-b, Extended Data Fig. 2**). The injection sites and the retrogradely-labeled neurons found inside the MLR locus defined in **Fig. 2** were registered to a common reference brain space^50^. Such retrograde tracing revealed neurons located in the MLR locus whose axonal projections descended both ipsi- and contra-laterally (**Fig. 4c1-2,** 11 larvae). We observed no difference in the number of cells labeled at either side of the injection site. To investigate the possible projection pattern of individual neurons located in the MLR region we employed “virtual single cell tracing” interrogating the cellular-resolution brain atlas of *mapzebrain*^50^. We searched for neurons with soma located in a region of 30 μm in radius medial and dorsal to the LC. Most of the neurons retrieved had ipsilateral descending axons to the reticular formation, with some of them projecting down to the most caudal medulla (**Fig. 4d,** see also **Extended Data Fig. 3**). Interestingly, we found that none of the axons reached the dorsal portions of the pontine and retropontine reticular formation.

**Fig. 4.**
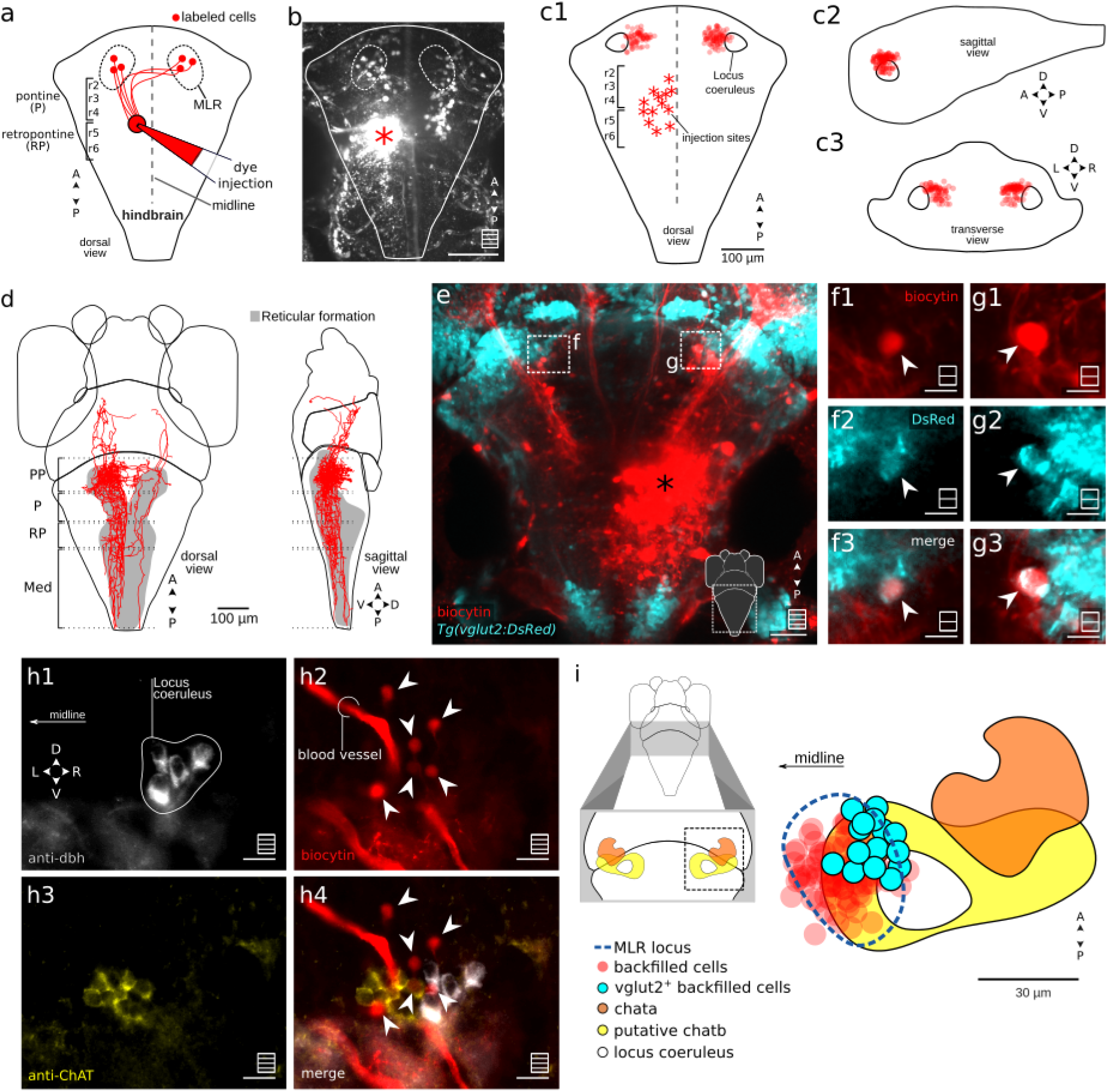
The MLR sends bilateral projections onto the reticular formation. **a,** Schematic representation of the retrograde tracing experimental design in which biocytin was unilaterally injected in the pontine and retropontine reticular formation between rhombomere 2 and 6 (r2-r6) to reveal neurons in the MLR locus projecting in the reticular formation (r, rhombomere). **b,** A typical Z projection stack acquired 2 hours after the backfill injection (asterisk) shows neurons with a soma located in the MLR area (dorsal view, white dash line circle, radius of ~30μm centered on the MLR). Scale bar is 100 μm. **c,** Location of all retrogradely-labeled neurons found in the MLR region denoted in panel **b** mapped onto a common brain space (11 fish, 156 cells, neurons backfilled contralateral to injection site: 8.00 ± 4.6, ipsilateral: 6.18 ± 4.6) in the dorsal **(c1)**, sagittal **(c2)** and transverse **(c3)** planes. The number of cells found ipsi-and contra-laterally to the injection site are not statistically different (one sample t-test; t(19.8) = 0.9 p = 0.35) Scale bar is 100 μm. **d,** 3D rendering of neurons with soma located in the MLR from the single-neuron atlas^50^ (15 neurons). **e,** Some of the retrogradely-labeled neurons were glutamatergic as shown by overlap with the *Tg(vglut2:DsRed)* transgenic line: a maximal Z-stack projection over 10 μm revealed retrogradely-labeled neurons (red) shown in the dorsal plane together with DsRed-positive cells (cyan). Biocytin injection is denoted with an asterisk. Scale bar is 25 μm. **f, g,** Detailed view of regions denoted in panel **e** and corresponding to single planes acquired at different depths. Arrowheads indicate putative glutamatergic retrogradely-labeled neurons. Scale bar is 5 μm. **h,** Photomicrograph illustrating an example of the distribution of labeled neurons in the MLR region as seen in epifluorescence microscopy showing somata that are immunoreactive to Dbh (white) **(h1)** and ChAT (yellow) **(h2)** in the vicinity of retrogradely-labeled MLR neurons (red) **(h3, h4). i,** Based on the *mapzebrain* zebrafish space^50^, schematic representation of the location of the retrogradely-labeled glutamatergic cells (cyan, 6 fish, 15 cells), *chata^+^* neurons (orange), *chatb*+ cells whose location was estimated from^56,57^ (yellow), locus coeruleus (white), and the MLR locus defined by the delimitation of stimulation sites that effectively triggered forward swimming (dashed line) with all neighboring retrogradely-labeled neurons (red). For panels **b**, **e, h**: maximum Z projection stacks (schematized in the bottom right corner with squares including multiple lines); **f**: single optical sections (schematized in the bottom right corner with squares including a single line). PP, prepontine; P, pontine; RP, retropontine; Med, medulla.

**Extended Data Fig. 2.**
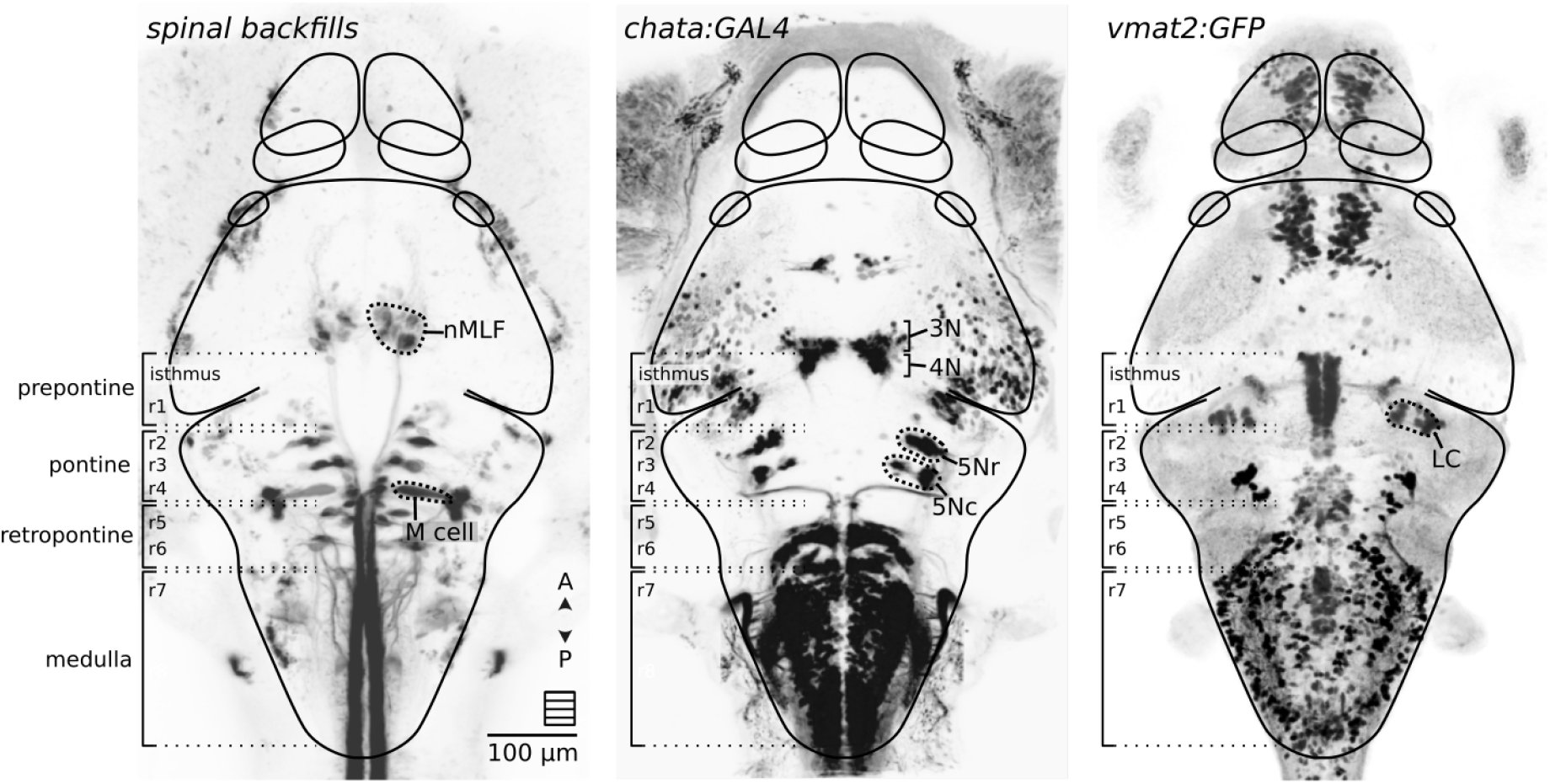
Brainstem nomenclature applied in this study according to^58^. The anatomical landmarks were defined based on the projecting neurons labeled by spinal backfills, the expression patterns of *Tg(chata:GAL4)* and *Tg(vmat2:GFP)* transgenic lines. All images were taken from the web-interface of *mapzebrain* (https://fishatlas.neuro.mpg.de/, download: 12 Apr. 2020). The *prepontine* region contains the isthmus and rhombomere 1 (r1). The trochlear nucleus (4N) lies in the rostral border of the isthmus and, amongst other structures, r1 contains the locus coeruleus (LC). The *pontine* region is defined rostrally by rhombomere 2 (r2), containing the rostral trigeminal motor nucleus (5Nr), the caudal part of the trigeminal motor nucleus (5Nc), and caudally by rhombomere 4 (r4) including the Mauthner cell (M cell). The *retropontine* region is composed of rhombomeres 5 and 6 (r5, r6) containing the abducens (6N) and facial nucleus (7N) respectively. The *medulla* is delimited rostrally by rhombomere 7 (r7) and caudally by rostral spinal cord. Under this nomenclature, the oculomotor nucleus (3N) is a midbrain structure, but the nucleus of the medial longitudinal fasciculus (nMLF) should be considered as a diencephalic structure.

**Extended Data Fig. 3.**
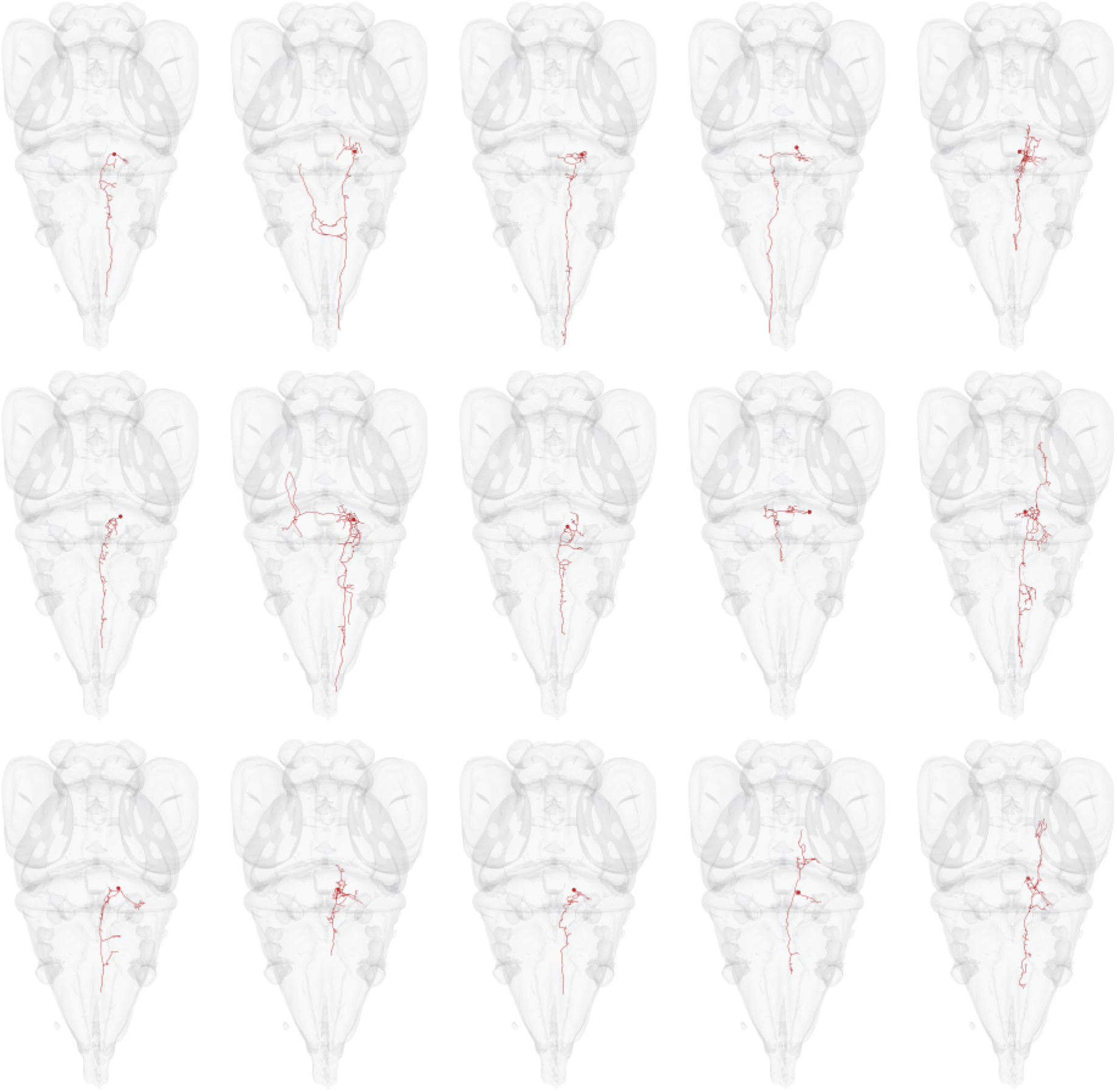
Individual 3D morphology of neurons shown in **Fig. 4d** whose soma is localized in the MLR. All images were taken from the web-interface of *mapzebrain* (https://fishatlas.neuro.mpg.de/, download: 12 Apr. 2020).

In lamprey as in mouse, the main synaptic output from the MLR that drives locomotion is glutamatergic^27,49,53^. To investigate whether some MLR neurons were glutamatergic in zebrafish, we performed backfill experiments targeting the reticular formation in the *Tg(vglut2:GFP)* transgenic line that labels numerous glutamatergic neurons^54^. We found that some of the retrogradely-labeled neurons in the MLR region were *vglut2*-positive (**Fig. 4e-g**). As glutamatergic neurons are intermingled with cholinergic neurons in the MLR of lampreys^49^, salamanders^17^ and mice^53,55^, we investigated in retrograde labeling experiments whether cholinergic neurons were located within the zebrafish MLR locus using immunohistochemistry against ChAT and Dbh. As seen on cross sections at the level of the MLR locus (see **Fig. 4c3**), retrogradely-labeled neurons were medial to Dbh-positive cells from the locus coeruleus and in close proximity to ChAT-positive cells, presumably belonging to the zebrafish homolog of the laterodorsal tegmental nucleus^29,29^ (LDT) (**Fig. 4h**). In agreement with this observation, mapping all the results obtained so far in the same reference brain space revealed that glutamatergic and cholinergic cells were in the vicinity of the loci whose stimulation elicited forward swimming (**Fig. 4i**). We found that the cholinergic cells expressing the *chata* gene in the tegmentum within the nucleus isthmi^56^ do not colocalize with the MLR, suggesting that the immunoreactive ChAT-positive neurons found centered on the MLR belong to the nucleus expressing *chatb* and *vachtb* genes^56,57^. Altogether, our anatomical investigations confirm that MLR neurons in zebrafish project onto the reticular formation and include glutamatergic cells in the vicinity of cholinergic neurons.

### MLR stimulation recruits hindbrain V2a RSNs in the pontine, retropontine, and medullary regions

We next sought to investigate whether the zebrafish MLR recruits V2a RSNs. Because V2a brainstem neurons are not all reticulospinal neurons^34,35,37,41,59^, we first quantified the proportions of V2a RSNs by optically-backfilling the V2a brainstem neurons projecting to the spinal cord by photoconverting their axons at the rostral spinal cord in 4.5 dpf *Tg(vsx2:Kaede)* transgenic larvae (**Fig. 5a1**, see **Methods**). We took confocal stacks of the hindbrain and observed that only 57% of the V2a neurons were RSNs in the dorsal pontine (dP: 57.5% ± 18.1%) and 35% of cells in the dorsal retropontine (dRP: 35.5% ± 6.1%) (**Fig. 5a2**). In contrast, a vast majority of V2a cells were RSNs in the ventral pontine nucleus (vP: 95.6% ± 5.1%, 4 fish), ventral retropontine (vRP: 84.1% ± 9.5%), and medulla (Med: 96.6% ± 1.8%).

Interestingly, the medulla contained 60% of all V2a RSNs. For our analysis on the recruitment of V2a RSNs upon MLR stimulation, only the ventral pontine, ventral retropontine and medullary regions were therefore included.

In lampreys and salamanders, a unilateral single pulse stimulation of the MLR elicits bilateral excitatory postsynaptic potentials in hindbrain RSNs^29,51,60^. In order to test whether the MLR drives ipsi- and contra-lateral RSNs in larval zebrafish, we investigated in *Tg(vsx2:GAL4;UAS:GCaMP6s)* transgenic larvae the response of V2a RSNs to unilateral single pulse (1-5 μA, 2 ms long) MLR stimulations (**Fig. 5b1-b2**). We found that unilateral stimulation of the MLR evoked bilateral excitatory responses in V2a neurons throughout the hindbrain from the pontine region to the medulla (**Extended Data Fig. 4a**).

We confirmed the specific MLR coupling to RSNs by analyzing the amplitude of the calcium transients in large identifiable ventral pontine and retropontine V2a RSNs named RoM2, RoM3, MiV1, MiD2i, MiD3i^37^ (**Fig. 5c1**). In agreement with previous reports^29,51,60^, these large ventrally located RSNs were reliably recruited starting at low stimulation intensity and similarly responded with an amplitude modulation scaling with stimulation intensity (**Fig. 5c2-c3**), further supporting our identification of the MLR in larval zebrafish.

Upon stimulation of graded intensities (**Fig. 5b2**), a subset of brainstem V2a neurons reliably responded above a given threshold (“reliably recruited neurons’“ in color, 22%, **Extended Data Fig. 4b, Extended Data Fig. 5a**), others either did not consistently respond (“unreliably recruited” in gray, 31%) or never responded (“non recruited”, 47%). We focused our analysis on reliably recruited cells as they likely receive excitatory inputs from the MLR and contribute to the motor output (5 fish, n = 391 out of 1781 recorded neurons). Half of V2a RSNs were reliably recruited in the ventral pontine and retropontine areas (**Fig. 5d**, 5 fish, vP: 42.8 %, vRP: 50.0 %, **Extended Data Fig. 5b**) and their recruitment occurred at the lowest stimulation intensity used (**Fig. 5e, Extended Data Fig. 4b, Extended Data Fig. 5c**). We investigated whether calcium responses of reliably recruited V2a RSNs exhibited different rising slopes upon MLR stimulation, as the firing rate of neurons correlates with the rising slope of their calcium transients^61,62^. Responses of V2a RSNs in the ventral pontine region and ventral retropontine (subset marked by arrowhead, **Fig. 5f1**) showed consistently large rising slopes, larger to the ones observed in any of the other regions (**Fig. 5f2, Extended Data Fig. 5d**), suggesting an effective and prompt coupling between MLR and ventrally located V2a RSNs.

In the medulla, a small proportion of V2a RSNs were reliably recruited (**Fig. 5e**, 5 fish, 17%). In contrast to RSNs in the pontine and retropontine areas, the recruitment of medullary V2a RSNs occurred in a graded manner as a function of MLR stimulation intensity (**Fig. 5e, Extended Data Fig. 4b**), suggesting that these cells could be involved in eliciting graded locomotion.

**Fig. 5.**
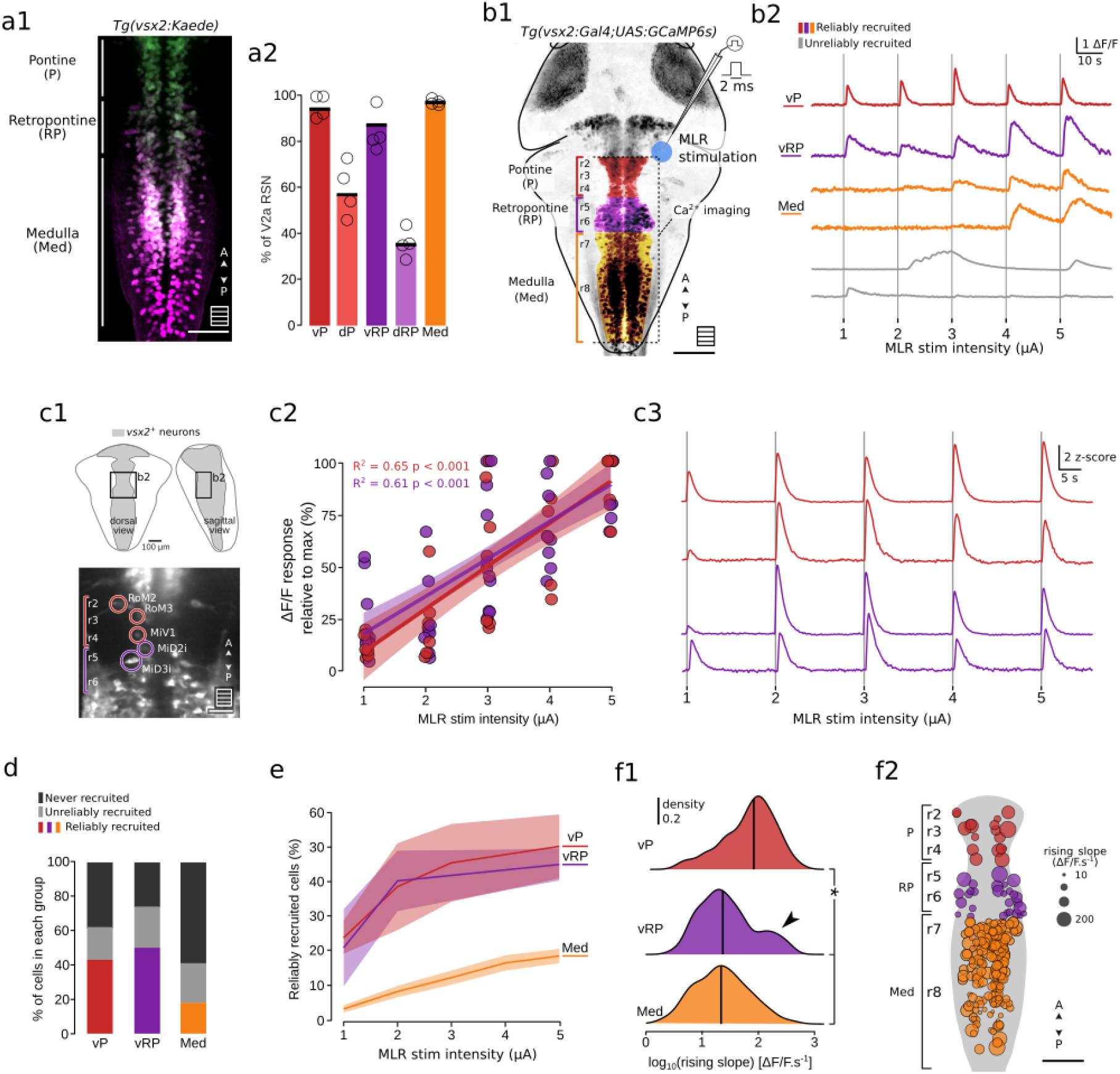
Single pulse MLR stimulations gradually recruit a subset of V2a RSNs. **a,** Distribution and density of RSNs within the V2a population estimated from optical backfills experiments. **a1,** Example image of the hindbrain of 5 dpf *Tg(vsx2:Kaede)* larval zebrafish after photoconverting Kaede in the rostral spinal cord at 4.5 dpf. While green cells do not project to the spinal cord, all magenta or white cells contain photoconverted Kaede and are therefore counted as RSNs. Scale bar is 50 μm **a2**, The proportion of V2a RSNs identified as projecting an axon to the spinal cord reveals that a vast majority of V2a neurons are RSNs in pontine and retropontine ventral areas as well as in the medulla. **b1,** Schematic illustration of the experiments investigating *vsx2+* (V2a) neurons recruitment in response to MLR stimulation. In paralyzed *Tg(vsx2:GAL4;UAS:GCaMP6s)* transgenic 6dpf larval zebrafish, local single 2ms-long 1-5 μA electrical current pulse MLR stimulations were applied using glass-coated tungsten microelectrode while the calcium activity of V2a RSNs was recorded in pontine, retropontine and medullary region (P, RP and Med respectively). **b2,** Typical calcium traces of reliably recruited V2a neurons in the different anatomical regions investigated, during a train of single shot MR stimulation of different intensities (top, colored traces: vP, ventral pontine; vRP, ventral retropontine; Med, medulla). Gray traces on the bottom represent neurons which exhibited a calcium response after at least one stimulation but were not labeled as reliably recruited. **c,** Identifiable V2a RSNs in the ventral pontine (vP)-retropontine (vRP) areas were reliably and gradually recruited by the MLR stimulation, **c1,** Example plane (top panel) showing the location of selected V2a RSNs (bottom panel, scale bar is 20um). **c2,** The amplitude of calcium responses of V2a RSNs in vP and vRP scaled with MLR stimulation intensity (3 fish, 1-2 planes per fish, 23 neurons, 2-9 neurons per plane). **c3,** Typical traces from identifiable V2a RSNs in the vP and vRP areas show fast rising slope and gradual response to increasing MLR stimulation intensity (1 typical fish, 2 planes represented). **d**, The recruitment of V2a neurons studied could be clustered into 3 groups: “reliably-recruited” (colored - color corresponds to reliably recruited neurons in a given anatomical area), “unreliably-recruited” (gray) and “non recruited” (black) (5 fish, total 1210 neurons; vP: 42 neurons, vRP: 88 neurons, Med: 1080 neurons) (see **Methods**). **e,** Proportions of reliably recruited V2a neurons responding at each MLR stimulation intensity (proportions are relative to numbers of V2a neurons in each anatomical group (defined in **b**). **f,** Kinematics of the V2a neurons’ calcium response to MLR stimulation. **f1,** Distribution of rising slopes of calcium transients in reliably recruited neurons shows larger values in the ventral pontine region compared to other regions. Distribution of rising slopes in other regions were indistinguishable from each other (5 fish, vP: 18 neurons, 76 events, 64.4 ± 67.6 ΔF/F.s^-1^; vRP: 44 neurons, 177 events, 30.6 ± 43.3 ΔF/F.s^-1^; Med: 183 neurons, 583 events, 24 ± 27.3 ΔF/F.s^-1^, Kruskal-Wallis test, χ^2^ = 141.68, p <0.001, Wilcoxon’s test pairwise comparisons: vP vs vRP: p < 0.01; vP vs Med: p < 0.001; vRP vs Med: p = 1.0). **f2,** Location of reliably recruited V2a neurons (dot size encodes the median of all computed rising slopes for each neuron).

**Extended Data Fig. 4.**
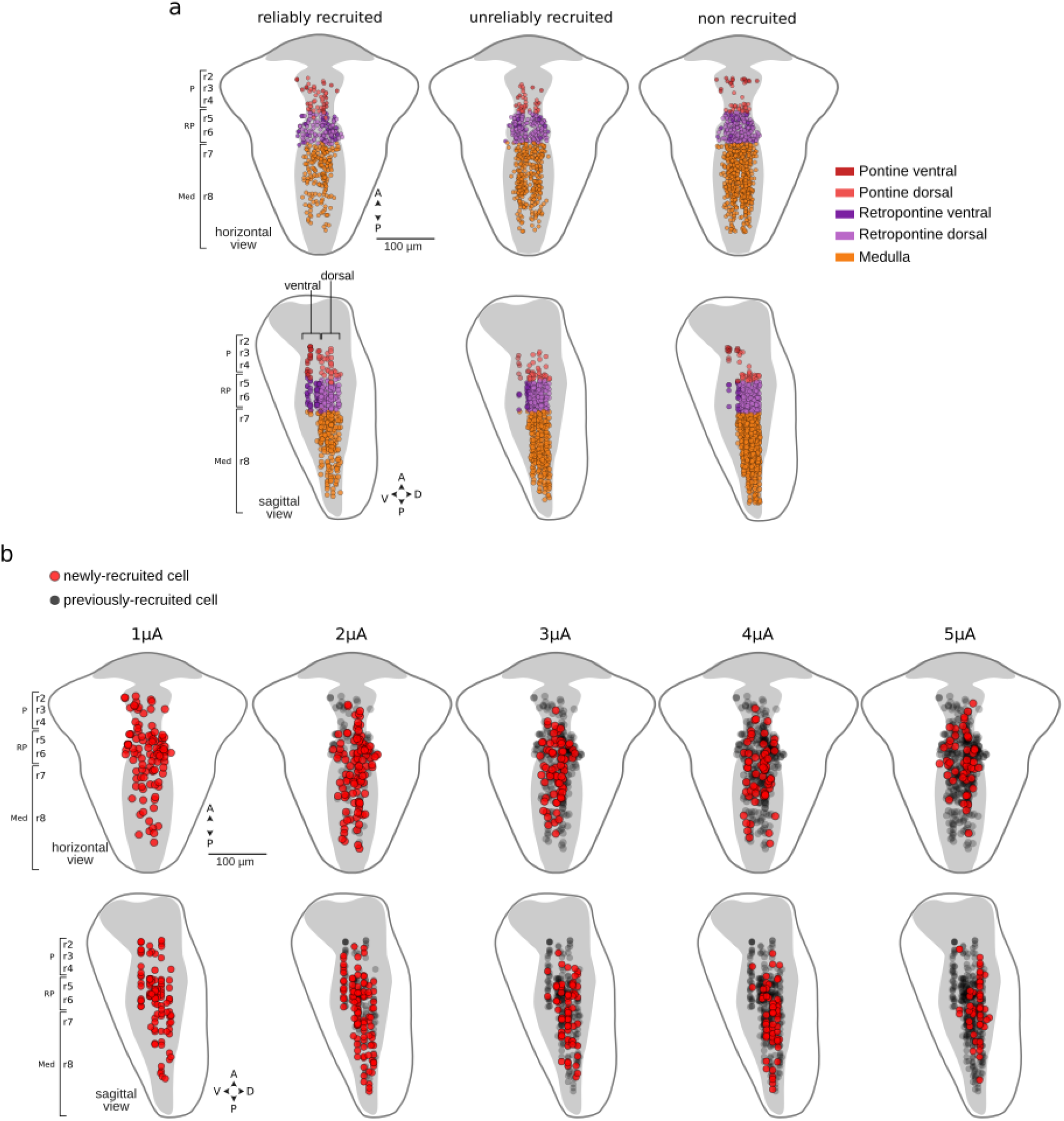
**a.** Location of V2a neurons reliably- and unreliably recruited and non-recruited throughout the pontine, retropontine and medulla. Horizontal (top) and sagittal (bottom) views are shown. **b.** Hindbrain location of the reliably recruited V2a neurons activated by MLR stimulation at increasing intensities (1 to 5μA). The red dots represent newly recruited neurons at a particular stimulation intensity. The gray dots show neurons previously recruited at lower intensity.

**Extended Data Fig. 5.**
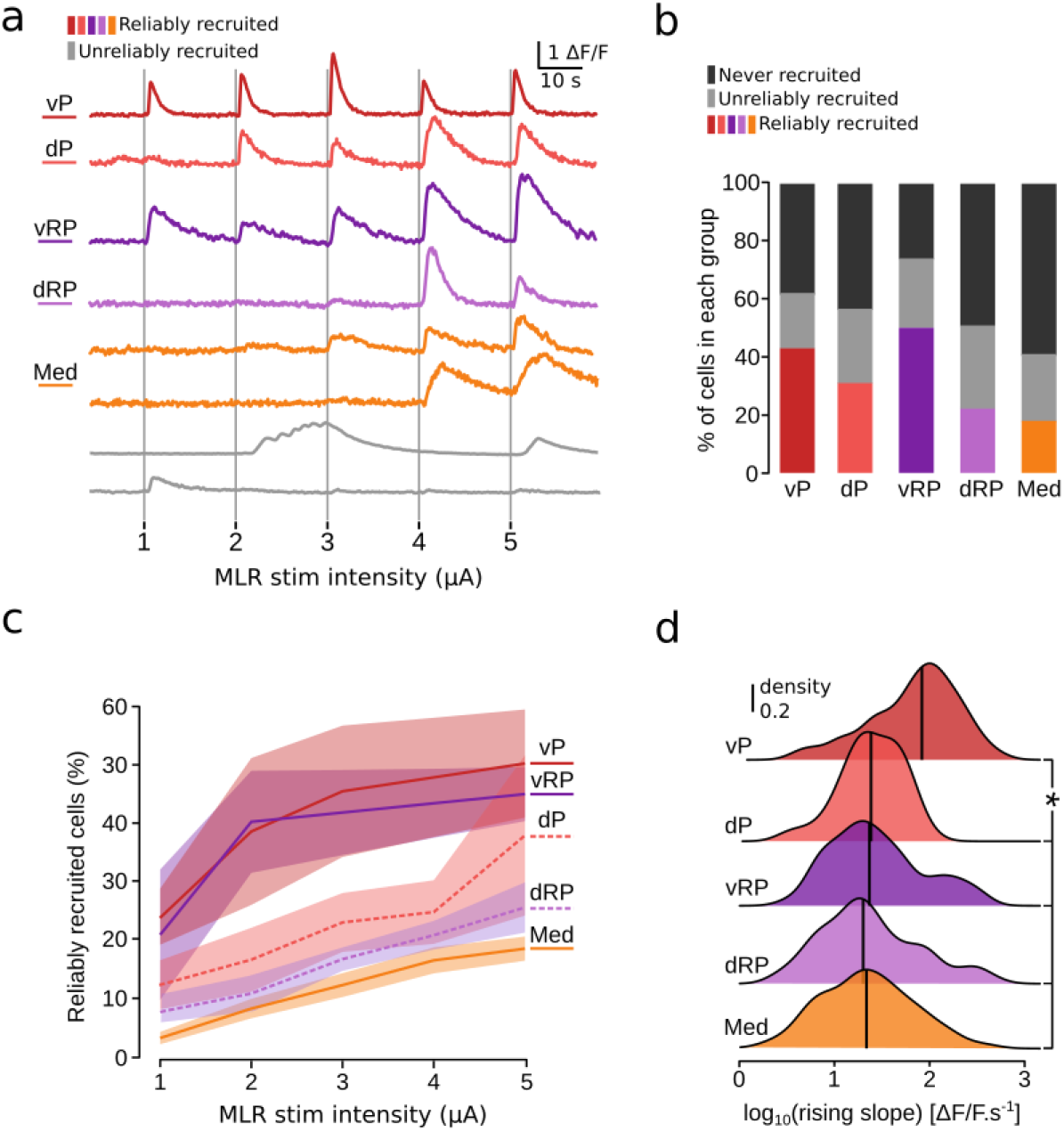
**a,** Typical calcium traces of reliably recruited V2a neurons in the different anatomical regions investigated (top, colored traces: vP, ventral pontine; dP, dorsal pontine; vRP, ventral retropontine; dRP, dorsal retropontine; Med, medulla). Gray traces on the bottom represent neurons which exhibited a calcium response after at least one stimulation but were not labeled as reliably recruited. **b**, The recruitment of V2a neurons studied could be clustered into 3 groups: “reliably-recruited” (colored - color corresponds to reliably recruited neurons in a given anatomical area), “unreliably-recruited” (gray) and “non recruited” (black) (5 fish, total 1781 neurons; vP: 42 neurons, dP: 96 neurons, vRP: 88 neurons, dRP: 475 neurons, Med: 1080 neurons). **c,** Proportions of reliably recruited V2a neurons responding at each MLR stimulation intensity. **d,** Distribution of rising slopes of calcium transients in reliably recruited neurons shows larger values in the ventral pontine region compared to other regions. Distribution of rising slopes in other regions were indistinguishable from each other (5 fish, vP: 18 neurons, 76 events, 64.4 ± 67.6 ΔF/F.s^-1^; dP: 29 neurons, 87 events, 23.4 ± 20.2 ΔF/F.s^-1^; vRP: 44 neurons, 177 events, 30.6 ± 43.3 ΔF/F.s^-1^; dRP: 100 neurons, 326 events, 27.1 ± 30.7 ΔF/F.s^-1^; Med: 183 neurons, 583 events, 24 ± 27.3 ΔF/F.s^-1^, Kruskal-Wallis test, χ^2^ = 141.68, p <0.001, Wilcoxon’s test pairwise comparisons: vP vs dP: p < 0.05; vP vs vRP: p < 0.01; vP vs dRP: p < 0.01; vP vs Med: p < 0.001; dP vs vRP: p = 1.0; dP vs dRP: p = 1.0; dP vs Med: p = 1.0; vRP vs dRP: p = 1.0; vRP vs Med: p = 1.0; dRP vs Med: p = 1.0).

### Subsets of V2a medial medullary neurons encode number of oscillations and locomotor frequency during forward locomotion

In order to test whether medullary V2a RSNs translate the MLR command into graded forward locomotion, we investigated whether these cells could encode locomotor frequency or amplitude of forward swimming. We performed functional imaging in head-fixed tail-free zebrafish larvae during long lasting forward episodes elicited by long MLR train stimulations (0.1 μA; 10 Hz; 40 s-long) (**Fig. 6a**). Such stimulation elicited unusually long symmetric forward episodes lasting up to 20 s and containing up to hundreds of oscillations (**Extended Data Fig. 6a-b**). We found that half of the medullary V2a neurons were active during locomotion (60.1 ±19 %, 133 ±37.2 out of 228 ± 35.9 neurons per fish, a total of 1140 neurons out of 5 fish recorded). As medullary V2a RSNs exhibited diverse patterns of activity, we clustered them based on their calcium signal (**Fig. 6b**, see **Methods**, **Supplementary Video 2**). We identified a cluster of medullary V2a RSNs that were specifically recruited during forward swims (**Fig. 6b,** referred to as “forward cluster”) and another specifically recruited during struggles (**Extended Data Fig. 6c**). Medullary V2a RSNs in the forward cluster were medially located (**Fig. 6c**) whereas the ones active during struggles were laterally located (**Extended Data Fig. 6d**). The proportion of V2a medullary neurons of the forward cluster was similar to the proportion of medullary neurons reliably recruited by the MLR (forward cluster: 18.32 ± 7.32%, 26.4 ±12.6 out of 228 ± 35.9 neurons per fish, a total of 1140 neurons out of 5 fish recorded; **Fig. 6,** reliably recruited V2a neurons: 22% of medullary V2a neurons).

We then focused our analysis on the activity of V2a RSNs associated with forward swimming. We noticed that the calcium signals of V2a RSNs in the forward cluster increased throughout the forward episode (**Fig. 6b**), suggesting that these V2a RSNs could encode the number of oscillations of forward swims. Accordingly, we found a strong correlation between the maximal amplitude of the calcium transient and the number of oscillations during a forward episode (**Fig. 6d1**) for numerous medial V2a RSNs dorsally located in the caudal medulla (**Fig. 6d2**). Using a regression-based approach to model the activity of these cells, we investigated which kinematic parameters the medullary V2a RSNs encode among tail beat frequency, amplitude of the tail beat and their modulation (see equations used to build the linear model on the calcium activity in **Methods**; **Fig. 6e, f1, f2**, **Extended Data 6d-e**). Notably, the activity of most medullary V2a RSNs in the forward cluster encoded the instantaneous tail beat frequency (**Fig. 6f1**, neurons whose coefficient was above 0 and significantly relevant: n = 59/89 neurons; see **Methods**). Interestingly, a group of V2a RSNs in the caudal and dorsal medulla both encoded the instantaneous tail beat frequency and the number of oscillations (**Fig. 6f2**). In contrast, the activity of other V2a RSNs in the ventromedial medulla encoded both instantaneous tail beat frequency and the rise in tail bend amplitude during forward swims (**Fig. 6f2**). Altogether, our results highlight that medullary V2a RSNs have a sustained activity during forward locomotion and mainly encode the instantaneous locomotor frequency.

**Fig. 6.**
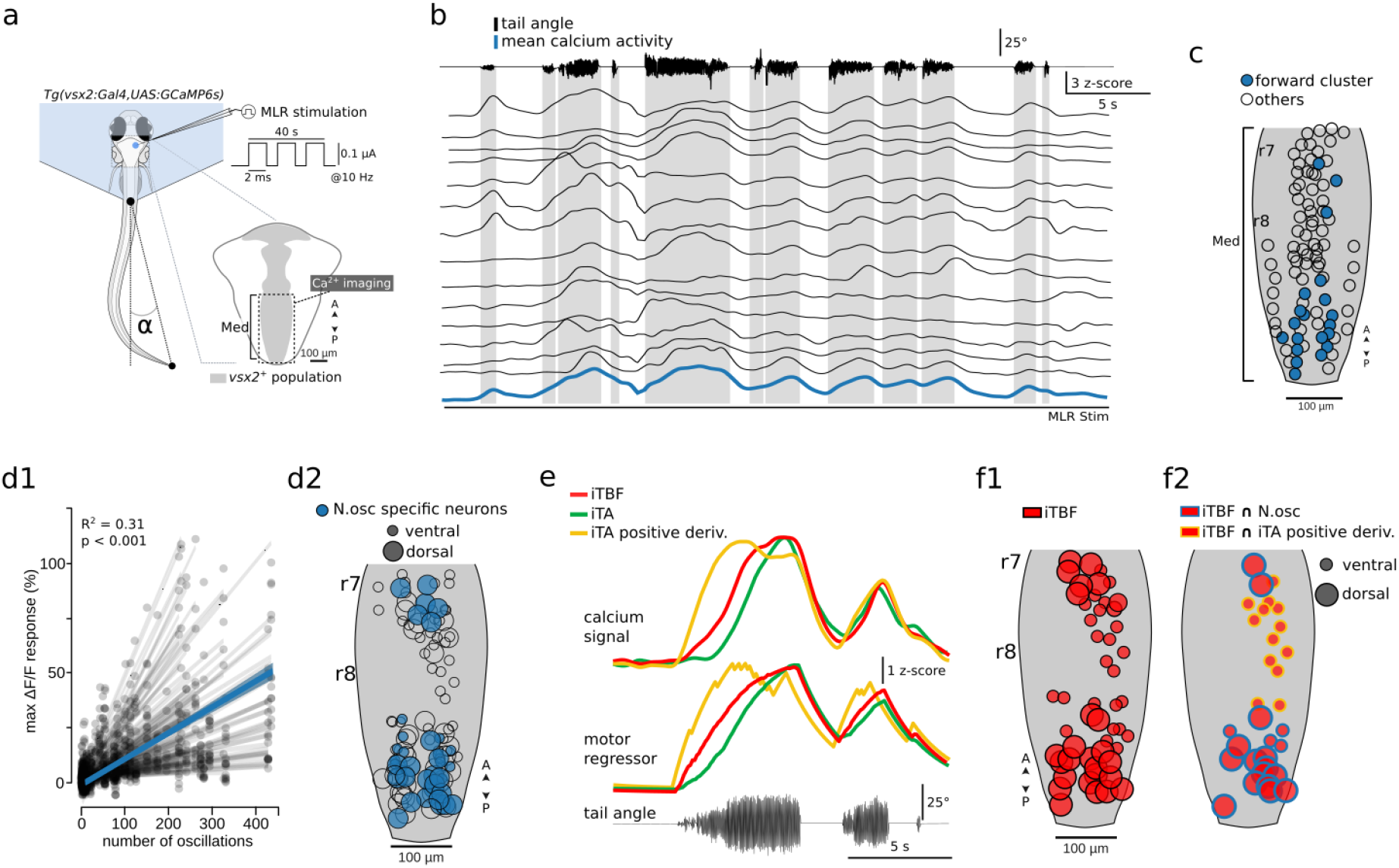
Medullary V2a RSNs are specifically recruited during MLR induced-forward swimming and encode the number of oscillations, tail beat frequency and amplitude. **a,** Schematic illustration of the experiment investigating medullary V2a RSN recruitment during MLR-induced forward swimming. **b,** Tail angle trace and calcium activity of a functional cluster of V2a RSNs active during forward swimming. For ease of visualization, only the tail angle for forward episodes is represented here. Mean neural activity of the group is plotted in blue, individual traces in the clusters in black (z-score). **c,** V2a RSNs whose activity is represented in (b) are in the medial medulla (filled circles in blue). Empty circles represent neurons that did not show any activity during the experiment or corresponded to different clusters than the forward one (see also Extended Data Fig. 4 c and d). **d,** The amplitude of the response of medullary V2a RSNs in the forward cluster correlated with the number of oscillations during a forward episode. **d1**, The plot shows a linear regression between the max ΔF/F of individual V2a RSN and the number of oscillations of the forward episodes (n = 132 neurons in n=5 fish, n = 8 trials). **d2,** Dorsal view showing the spatial distribution of neurons whose response amplitude correlated the most with the number of oscillations (blue filled circles, n =38/132 neurons in n = 5 fish, n = 6/8 trials, p < 0.12 and correlation coefficient >= 0.8). Empty circles correspond to other V2a RSNs from the forward cluster whose response amplitude was less correlated with the number of oscillations during a forward episode. **e.** Individual neurons’ activity reflect 3 kinematic parameters: tail beat frequency, tail beat amplitude and binary positive derivative of the tail amplitude. *Top:* traces from 3 example neurons that were recruited during forward episodes and whose calcium activity differed during the episode *Middle:* 3 motor regressors that best recapitulated the calcium activity of the 3 neurons above (matching color codes) with high correlation for instantaneous tail beat frequency (red), instantaneous tail amplitude (green) or the increase in tail amplitude (yellow). *Bottom* corresponding tail angle trace. **f1,** Subsets of V2a medullary RSNs encode the instantaneous tail beat frequency. The circle size encodes their location along the dorsoventral axis (n=5 fish, n=6 experiments, n= 59/89 neurons); **f2**, dorsal view showing the spatial distribution of V2a RSNs whose activity encoded tail beat frequency and either increase in tail beat amplitude (yellow outline) or number of oscillations (blue outline, from panel d2).

**Extended Data Fig. 6.**
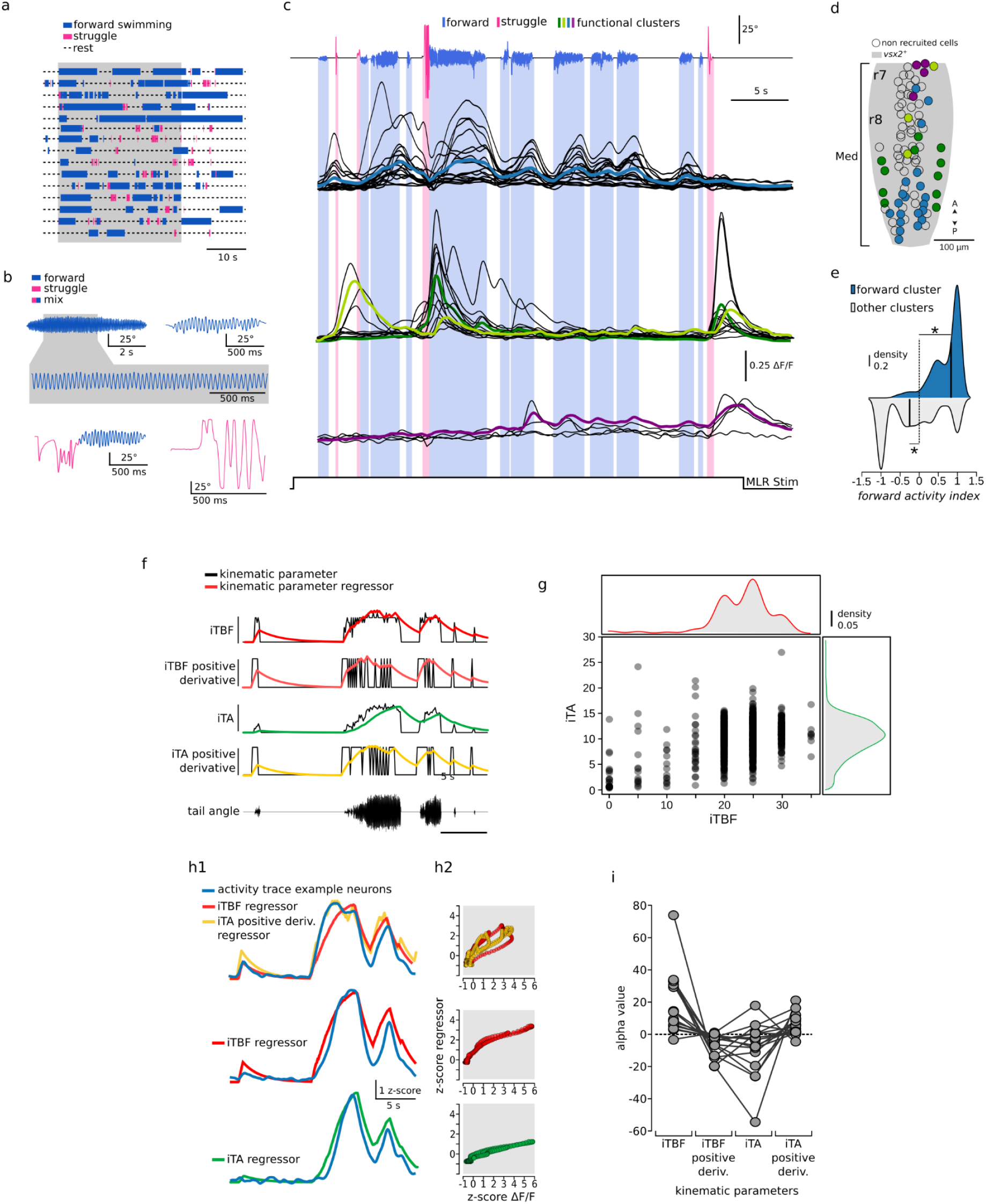
**a.** Schematic representation of behavioral traces during all experiments investigated in Fig 6. Episodes are represented as filled boxes whose color depict the behavior type (blue: forward episode, pink: struggle). **b.** Example bouts elicited during 40s long MLR stimulation corresponding to either pure forward swimming (top trace), bout of mixed episodes (bottom left) or pure struggle. Episodes are color-coded by manually curated categories. **c.** Tail angle trace and calcium activity clustered by similarity during the same example experiment than **Fig. 6b**. Here all functional clusters are represented. Each behavioral episode is color-coded by manually curated categories (blue: forward swimming, magenta: struggle behavior). Mean neural activity for each cluster is represented in color-trace and individual traces in the clusters in black. Neurons in blue are the forward cluster shown in Fig. 6b. Neurons in the green clusters are active during struggle episodes. **d.** Dorsal view showing the spatial distribution of neurons color-coded by the functional cluster defined in **c**. Neurons in the green clusters are laterally located. Non-filled dots represent neurons recorded which did not show any activity during the experiment. **e.** Medullary V2a RSNs in the forward clusters were more recruited during forward swimming than during struggle behavior. *Forward activity index* calculated from the calcium activity as (*forward activity index* = (average (max ΔF/F during forward episodes) - average (max ΔF/F during struggle) / average (max ΔF/F during forward episodes) + average (max ΔF/F during struggle)). A positive index indicates an average max ΔF/F higher during forward than during struggle behavior (5 fish; forward cluster: 1438 cells (mean ± sd): 0.71 ± 0.37, one sample t-test against mu = 0; t(1437) = 72 p < 0.001; rest of the clusters: 2603 cells: −0.17 ± 0.72, one sample t-test against mu = 0, t(6626)=-12.6, p < 0.001). **f,** Mean activity of an example forward cluster and the 4 motor regressors encoding distinct kinematic parameters: the tail beat frequency, the tail bending amplitude, and their binary positive derivative. Each motor regressor simulates the calcium trace of a neuron whose spiking activity would perfectly recapitulate a motor feature. Top trace, mean activity of an example forward cluster. Middle traces, raw signal (black) and corresponding regressor (color) of each motor feature represented. Bottom trace, corresponding tail angle. **g.** Distribution and correlation between the different instantaneous swimming parameters. Instantaneous tail beat frequency and tail amplitude are co-dependent, but not linearly correlated. Distribution of the computed instantaneous tail beat frequency (red, top) and instantaneous tail amplitude (green, right) (2176 time steps, in 4 fish, 8 trials). **h,** Single neurons are explained by different motor regressors. **h1,** Neural activity of single neurons shown in **Fig. 6e** overlaid with the motor regressor that best explains it. **h2,** At each time step, the value of the motor regressor (x axis) is represented as a function of the mean activity of the forward cluster (y axis). **i,** Correlation coefficient of each motor regressor tested in the multiple linear model and illustrated for one example plane in one fish.

## Discussion

Our study provides the first identification of the locus of the zebrafish MLR in the prepontine region of the hindbrain. We found that electrical stimulation of the MLR induces sustained forward locomotion lasting for the duration of the stimulation and mimicking bouts deployed by larval zebrafish during spontaneous navigation. Here, we leveraged the genetic toolkit of larval zebrafish to investigate the functional coupling of the MLR to a key population of RSNs, the V2a RSNs, labeled by the transcription factor *vsx2*. We show that the MLR sequentially recruits a subset of V2a RSNs throughout the brainstem: while pontine and retropontine V2a RSNs are recruited at once at low intensity, medullary V2a RSNs are progressively recruited upon increasing intensity of the MLR stimulation. By taking advantage of the exceptionally long forward swims induced by MLR stimulation, we were able to uncover a previously overlooked group of medullary V2a reticulospinal neurons whose activity is maintained during forward locomotion. In contrast to lateral medullary V2a RSNs recruited during struggles, medial V2a RSNs in the medulla encode essential parameters determining the duration and speed of forward locomotion: the number of oscillations, the locomotor frequency and the increase in movement amplitude.

### Identification of the MLR in larval zebrafish

In all vertebrate species, the MLR sits at the mesopontine border along the cholinergic nuclei spanning from the caudal border of the substantia nigra to the locus coeruleus^16,17,53,63,64^. In mammals, the MLR comprises the pedunculopontine nucleus (PPN) and cuneiform nucleus (CuN) in the rostral mesopontine cholinergic stripe^15,19,24^. In salamanders and lampreys, the MLR is located in the caudal mesopontine cholinergic stripe corresponding to the laterodorsal tegmental nucleus^16,17^ (LDT). Here, we identified, based on electrical stimulations and anatomical investigations, the MLR of larval zebrafish as a small region of approximately ~30 μm in radius, which is medial and dorsal to the locus coeruleus and extends rostrocaudally in the prepontine area. Similarly to the location found in lamprey and salamanders, the zebrafish MLR colocalizes with cholinergic cells in the isthmus^56,57^, likely corresponding to the LDT^49^.

Even though stimulation in the region of the nMLF in fish induces locomotion^42–46,65,66^, the nMLF does not correspond to the MLR since it is not located within the mesopontine border^67^. The locomotor output elicited by nMLF stimulation is consistent with the motor responses observed upon activation of reticulospinal neurons^26,68–70^. Instead the nMLF should be considered primarily as a sensorimotor coordination center^71^ since it receives sensory inputs from the visual^72,73^ and vestibular systems^74^ and in turn sends descending projections^67,75^ to control head-tail positioning^66^ and spinal motor neurons^76,77^. The activity of the nMLF correlates with spontaneous^66,71^ and sensory induced locomotion^65,66,71,78^, probably due to the fact that eyes and tail movements are tightly coupled^79^. In zebrafish, the term “nMLF” was adopted in the first descriptions of the spinal projecting neurons^67,80^. Nevertheless, the nMLF corresponds to a structure called across vertebrate species the interstitial nucleus of Cajal (INC) (for review, see^81^). Indeed, the INC projects to the spinal cord^82^ and coordinates eye and head movements assisting the body in adjusting posture^83,84^ (for review, see^85^). Furthermore, INC stimulation can elicit locomotion in cats and rats^86^. Based on its anatomical and functional similarities, we propose that the nMLF should be renamed as the INC, as formulated for other vertebrate species (for review, see^87,88^).

### Control of locomotor output

Animals move using different types of gaits. Quadrupedal mammals walk at low speeds, trot at intermediate speeds and gallop at high speeds (for review, see^89^). The MLR controls the locomotor speed and its stimulation at different strength evokes the different gaits naturally exhibited by a given animal (for review, see^90^). For instance, stimulating the MLR in salamanders at low intensity produces a terrestrial stepping (walking), but the animal transitions to swimming (a faster gait) at higher stimulation strength^17^. In larval zebrafish, evidence for two different forward gaits have been reported in freely swimming conditions during optomotor response at different grating speeds^65^ (~27 and ~34 Hz each). Upon MLR stimulation we could elicit two clear locomotor regimes, one at 17 Hz for MLR stimulation below 15 Hz, and another at 23 Hz for stimulation at 20 Hz. We propose that these two locomotor regimes elicited correspond to the adaptation, in the head-embedded condition, of the two gaits described in freely swimming larval zebrafish. Larval zebrafish swims in a beat-and-glide fashion, where short periods of activity of typically few hundreds of milliseconds^91,92^ are followed by long interbout intervals where the animal is not actively swimming^93^. One of the most notable findings of this work is that by stimulating the MLR we were able to drastically prolong the duration of their swimming bouts, reaching up to 20 seconds.

### Functional coupling of the mesencephalic locomotor region to V2a reticulospinal neurons

In line with previous studies identifying the MLR (for review, see^24,94^), we demonstrate using retrograde tracing that the zebrafish MLR neurons send descending projections to the reticular formation. Experiments done in lampreys, mice and cats showed that only a small proportion of RSNs are active during spontaneous or MLR induced forward locomotion (lampreys^10^, cats:^95^, mice^35^). In agreement with these observations, we found that the MLR recruited a low proportion of brainstem V2a RSNs. More specifically, we show that the MLR recruits ventral V2a RSNs in the pontine and retropontine regions in an all-or-none fashion, consistent with large excitatory postsynaptic potentials previously recorded in these cells at low MLR stimulation intensity^29,51,60^. In contrast, the MLR gradually recruited, upon increasing intensities of stimulation, medial V2a RSNs in the medulla, similarly to what had been observed in lampreys and salamanders in unidentified RSNs^51,60^ that may include V2a neurons.

### Medullary V2a RSNs act as maintain cells encoding the instantaneous locomotor frequency

We find that a subset of medial medullary V2a RSNs sustain their activity during long episodes of forward locomotion elicited by MLR stimulation. In zebrafish, the recruitment of the large V2a RSNs in the ventral pontine and retropontine region had been well documented^37,40,78,96^. Yet, less was known on the role of medullary V2a RSNs found active during brief episodes of spontaneous fictive locomotion^59^. Here, we show that the calcium activity of a subset of V2a RSNs located in the caudal medulla scales with the number of oscillations within a bout, suggesting that this subset of V2a RSNs could operate as “maintain V2a RSNs”. Consistently, the ablation of medial V2a RSNs in the medulla in larval zebrafish resulted in shorter forward bouts^59^. Zebrafish medullary V2a RSNs project their descending axon ipsilaterally^36^ and resemble medullary RSNs previously recorded in xenopus tadpoles^97^, where they have been found to spike in phase and preceding each motor burst. Remarkably, zebrafish “maintain” V2a RSNs we highlight here are located in rhombomere 8, in the caudal most medulla at the boundary with the spinal cord. This region in lamprey has been found to generate the locomotor rhythm in isolation^98^. Multiple evidence indicate that this area may act as a critical hub to gate information from brainstem to spinal cord as its targeted electrical and optogenetic stimulation robustly induced locomotion^99–101^. By modeling the neuronal calcium activity with motor regressors for amplitude and frequency modulation, we uncovered that the activity of most medial V2a RSNs in the medulla encodes the locomotor frequency and for a subset, the increase in movement amplitude. Altogether these findings indicate that medial V2a RSNs in the medulla act as maintain cells setting the speed of forward motion, likely by recruiting spinal CPG interneurons controlling locomotor frequency^102,103^ and motoneurons to increase movement amplitude^59^. The previously overlooked V2a medullary RSNs we unveiled here may act in concert with other glutamatergic RSNs in the LPGi nuclei previously involved in mice in the control of locomotor speed^26^.

Here, we identified the MLR in larval zebrafish and mapped the downstream command circuits involved in forward locomotion. Altogether, this study represents a breakthrough for future investigations of supraspinal motor control, as virtually each and all reticulospinal neurons can be identified in the brain atlas, recorded and manipulated during active locomotion in this transparent and genetic model organism.

## Code availability

All python scripts used to process the data will be available upon publication of the manuscript at https://github.com/mathildelpx/Carbo-Tano_Lapoix_2022.

## Acknowledgements

We thank Sophie Nunes Figueiredo, Monica Dicu and Antoine Arneau from Animalliance for fish care. We thank Tod Thiele and Minoru Koyama for their comments on the results. We are grateful to Harold Burgess and Ashwin Bhandiwad for providing critical inputs to the manuscript. We thank Faustine Ginoux for the morphological registration pipeline and Gautam Sridhar for his inputs on the modeling approach. We thank Danielle Veilleux for making the monopolar micro-electrodes and her help with the histology.

## Contributions

M.C.T performed experiments, data extraction, analysis, plotting and statistical testing. M.L performed data extraction, analysis, mining and formatting, plotting, computations and registration. X.J performed optical backfills experiments, F.A performed immunohistochemistry and guided the anatomical investigations. M.C.T edited all figures. M.C.T and M.L drafted the first version of the manuscript. M.C.T, M.L, C.W wrote the manuscript with input from all authors. R.D and C.W conceptualized the study.

## Methods

### Animal care and transgenic lines utilized

Animal handling and procedures were validated by the Paris Brain Institute (ICM) and the French National Ethics Committee (Comité National de Réflexion Éthique sur l’Expérimentation Animale; APAFIS # 2018071217081175) in agreement with EU legislation. To avoid pigmentation, all experiments were performed on Danio *rerio* larvae of AB background with *mitfa (-/-)* mutation^104,105^. Adult zebrafish were reared at a maximal density of 8 animals per liter in a 14/10 hr light/dark cycle environment at 28.5°C. Larval zebrafish were typically raised in petri dishes filled with system water following the same conditions in terms of temperature and lighting as for adults. Transgenic larvae expressing Kaede and used for photoconversion experiments were exceptionally raised in complete darkness to minimize global photoconversion at early stages. Experiments were performed at 20°C on animals aged between 4 and 7 days post fertilization (dpf) as described in each experimental protocol below.

The following transgenic lines were used in this study: *Tg(vmat2:GFP)*^106^; TgBAC(vsx2:Gal4)^37^, TgBAC(vsx2:Kaede)^107^, Tg(UAS:GCaMP6s) (Muto et al., 2017)^66^; Tg(vglut2a:lRl-GFP)^108^.

### MLR electrical stimulation

Homemade glass-coated tungsten microelectrodes (0.7-3.1 MΩ, 10-35 μm exposed tip) were used to perform monopolar electrical stimulations. The electrodes positioned with a 35° angle penetrated the skin at different sites in the prepontine region using a motorized micromanipulator (MP-285A, Sutter Instruments, California, USA). In order to guide the electrode position to the MLR, we used the locus coeruleus as a landmark by labeling it with GFP in the *Tg(vmat2:GFP)^zf710Tg^* transgenic larvae expressing GFP under the *vmat2* promoter^106^. Stimulation was delivered using an isolated stimulation unit (2100, AM systems, USA) controlled with a Digidata series 1440A Digitizer (Axon Instruments, Molecular Devices, USA) to synchronize the electrical stimulation and behavioral recording. The timing of the electrical stimulations was triggered and recorded using Clampex 10.3 software (Axon Instruments, Molecular Devices, USA). When searching for the MLR, the behavioral response to stimulations in the locus of the MLR was typically recorded for multiple loci by applying continuous stimulation at 10 Hz and 1 μA not exceeding 30 s. If after 5 loci tested, no clear behavioral response was observed, the larva was discarded. To ensure success, a rest period of at least 3 min was allowed between successive trials. Once the proper site for the MLR was found, we implemented experimental trials lasting each about one minute. The stimulation protocol consisted of negative pulses of 2 ms duration, 5-20 Hz frequency at 0.1-3 μA intensity for 1-4s. To allow recovery and avoid synaptic habituation, the resting period between trials was at least 2 minutes. For experiments on paralyzed *Tg(vsx2:GAL4; UAS:GCaMP6s)* transgenic zebrafish, the stimulation protocol consisted of single 2 ms-long negative pulses with increasing intensities (1 to 5 μA) with steps of 1 μA and inter trial intervals of 20 s. For the same fish, a rest period of at least 3 min was allowed between trials performed at different imaging planes. All electrical stimulation experiments were carried out in a confocal microscope combining an upright microscope (Examiner Z1, Zeiss, Germany), using a 20X/1.0 DIC D = 0.17 M27 75mm (Zeiss, no:421452-9880-000) objective, a spinning disk head (CSU-X1, Yokogawa, Japan) and a laser light source (LaserStack, 3i Intelligent Imaging Innovations, USA).

### Behavior recording and analysis

#### Behavioral recording

6-7 dpf transgenic *Tg(vmat2:GFP)* or *Tg(vsx2:GAL4;UAS:GCaMP6s)* larvae were embedded in 2.5% low melting point agarose (Invitrogen, Thermo Fisher Scientific, USA) in a 35 mm Petri dish. The dish was then filled with external bath solution (134 mM NaCl, 2.9 mM KCl, 2.1 mM CaCl2-H2O, 1.2 mM MgCl2, 10 mM glucose, and 10 mM HEPES, with the pH adjusted to 7.4 and osmolarity to 290 mOsm). The agarose was removed caudally to the swim bladder in order to leave the tail free to move. Head-embedded larval zebrafish were illuminated from the side with a 45° angle using an 890 nm LED (ILH-IW01-85SL-SC211-WIR200, Intelligent LED Solutions, Berkshire, UK) and behavior was recorded at 300 Hz from below through a 5X20X/0.25 microscope objective 12.5mm (Zeiss, no:440125-0000-000) and imaged on a high-speed camera (acA640-750um, Basler, Ahrensburg, Germany) with a 50 mm lens (MVL50M23, Navitar, New York, USA). When the behavioral recording was coupled with functional calcium imaging, the 488 nm imaging laser was blocked with a 600 nm high pass filter located in front of the camera. The high-speed camera was controlled using the Hiris software (RD Vision, https://www.rd-vision.com/r-d-vision-eng, Saint-Maur-des-Fossés, France).

#### Tail tracking and automated segmentation of bouts

We used the open-source software ZebraZoom (https://github.com/oliviermirat/ZebraZoom) to track the position of the tail for each frame and extract bouts defined as discrete events when the tail was continuously moving^91,109,110^. The tail angle was defined as the angle between the body axis of the fish and the tip of the tail^91^. The quality of the tail tracking was carefully assessed by visual inspection. Electrically-induced bouts were defined as all bouts starting *during* the electrical train stimulation. Swim bouts starting just before the stimulation were excluded from analysis. “Spontaneous” bouts were defined as bouts starting and ending *before* the first electrical stimulation of the trial. Note that bouts occurring in the 1 min long time window *after* a stimulation train were excluded from the analysis as their properties could have been impacted by the prior electrical stimulation.

#### Segmentation of episodes within a bout

In the behavioral responses to electrical stimulations, bouts sometimes occurred as episodes of forward swims and struggles (see **Fig. 1, 2, 6**). We therefore manually segmented all bouts into episodes of either forward swims or struggles. The procedure was blindly implemented by an investigator who ignored the electrode position and the stimulation protocol used. Three rules were followed to maximize consistency: *i)* forward swims were defined by symmetric tail angle traces below 25 degrees with a minimum of three oscillations; *ii)* long forward episodes with rare tail bends above 25 degrees were kept as single forward episodes; *iii)* struggle episodes were defined as series of very large bends (tail angle > 25°) assigned as one single episode, but segmented into multiple episodes when bends were separated by more than 100 ms.

#### Analysis of kinematics

From the tail angle, the ZebraZoom algorithm extracted the timing and amplitude of each tail bend that we then used to calculate instantaneous tail beat frequency and instantaneous tail bend amplitude^91,109,110^. For each episode (**Fig. 1c, 2**), we estimated the corresponding duration, number of oscillations, and maximal tail bending amplitude as previously done for bouts in freely-swimming animals^91,109,110^, together with median bend amplitude and median instantaneous tail beat frequency (https://github.com/mathildelpx/Carbo-Tano_Lapoix_2022/).

#### Behavioral forward index

In order to estimate the efficiency of a stimulation site to elicit forward swimming, we computed for each stimulation trial a forward index defined as:

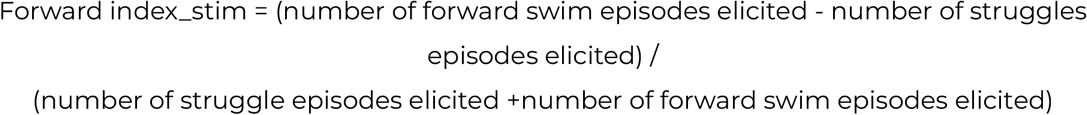

As we describe in the results section and in Extended Data Fig. 1, electrical stimulation at frequencies lower or above 10 Hz, or with intensities above 2 μA elicited mainly struggle behaviors. For this reason, we computed the median forward index of each stimulation site (Fig. 1e) only using the stimulation trials with optimal stimulation parameters, i.e.trains of pulses at 10Hz with current amplitudes of 1-2uA.

### Anatomy

#### Registration of electrical stimulation sites in the brain atlas

To assess the position of the stimulation electrode, Z projection confocal stacks were acquired combining the GFP signal and the differential interference contrast (DIC) channel using a 20X/1.0 DIC D = 0.17 M27 75mm (Zeiss, no:421452-9880-000) objective. The z-stacks were acquired with a step of 1 μm depth using Slidebook software 6.0 (3i Intelligent Imaging Innovations, Inc., Denver, CO, USA). From each image volume, we annotated manually in Fiji^111^ (http://fiji.sc/Fijji) the coordinates of the electrode position from the DIC channel and three anatomical reference coordinates from the GFP pattern of expression of the *Tg(vmat2:GFP)* transgenic line (medial boundary of the locus coeruleus on each side, anterior limit of the superior raphe and anterior limit of the inferior raphe). Using a custom-written python script, the coordinates of the anatomical landmarks were transformed to the mapzebrain fish space for the *Tg(vmat2:GFP)* transgenic line (https://fishatlas.neuro.mpg.de/, download: 12 Apr. 2020) and the resulting transformation matrix was then applied to the electrode coordinates.

#### Retrograde-labelling experiments using biocytin

6 dpf larvae were anesthetized with 0.02% MS-222 (A5040, Sigma-Aldrich) and embedded dorsally in 1.5% low melting point agarose prepared in external solution (see above). A small portion of the agarose was removed on top of the hindbrain and all the liquid was then extracted from the dish. A few crystals of biocytin Alexa Fluor 594 (Thermo Fisher, A12922) were diluted in 5 μL of dH2O, and then allowed to recrystallized on the tip of insect pins (stainless steel Minutiens, 0.2 mm diameter, Austerlitz). Under a stereo dissecting microscope, a unilateral single lesion in the hindbrain between rhombomere 2 and 6 was done using the dye-soaked insect pins. After the procedure, the dish was filled with external solution, the larvae were unmounted and allowed to recover for 2 hrs in oxygenated external solution. After larvae were again anesthetized with 0.02% MS-222 and mounted in 1.5% low melting point agarose. The retrogradely-labeled neurons were then imaged in a confocal microscope as described above. From each image volume, the coordinates of the backfilled neurons in the region of the MLR were annotated using Fiji. The positive backfilled neurons selected for this analysis were taken from the MLR area defined by electrical stimulations (with a diameter of approximately ~30 μm). For the experiment in Fig. 3a-d, the injections were done in WT fish. Since the dye injection forms a background labeling in the entire nervous system, we chose four anatomical landmarks also easily distinguishable in the mapzebrain *DAPI* fish space. For the experiments in Fig. 3e-g the four anatomical landmarks were selected on the basis of the DsRed fluorescence pattern in each transgenic *Tg(vglut2:DsRed-GFP)* larva (referred to as *Tg(vglut2:lox-DsRed-lox-GFP))* and from the *mapzebrain Tg(vglut2a:DsRed)* fish space. The script used and the registration steps applied were the same as described in the precedent section.

#### Immunohistochemistry

6 dpf larvae were processed as described above in order to label MLR neurons projecting to the reticular formation. After the injection, the larvae were allowed to recover for 2 hrs in oxygenated external solution. The larvae were then euthanized using 0.2% MS-222 and fixed in PFA 4% in PBS for 2 hrs at room temperature and then washed 3 times during 30 min in 1X PBS. The whole larvae were then transferred into a solution of 20% sucrose overnight at 4 °C. The next day, they were quickly frozen in methylbutane cooled to −45 °C and the heads were cut in a cryostat at 25 μm thickness in the transverse plane. The sections were collected in phosphate buffered saline containing 0.1% Triton X-100 (PBST, 0.1 M, pH 7.4, 0.9 % NaCl) and then blocked with PBST containing 10 % normal donkey serum (017-000-121, Jackson Immunoresearch, West Grove, PA) for 60 min at room temperature. The sections were then incubated for 48 h at 4 °C in PBST containing the following primary antibodies: Rabbit anti-dopamine-beta-hydroxylase (RRID: AB_572229, cat. number 22806, lot: 1905001, diluted 1:400, Immunostar, Hudson, WI) and goat anti-choline acetyltransferase (RRID: AB_2079751, cat. number AB144P, lot: 3430597, diluted 1:100, Millipore, Burlington, MA). The sections were then rinsed three times 10 min in PBST and incubated for 4 h in PBST containing the following secondary antibodies: Donkey anti-rabbit conjugated to Alexa Fluor 350 (cat. number A10039, diluted 1:200, Invitrogen, Eugene, OR) and donkey anti-goat conjugated to Alexa Fluor 594 (cat. number A11058, diluted 1:400, Invitrogen). The sections were then rinsed three times 10 min in PBS, mounted on ColorFrost Plus microscope slides (Fisher Scientific, Pittsburgh, PA) and coverslipped using Vectashield (H-1000, Vector Laboratories, Burlingame, CA) as mounting medium. The sections were then observed and photographed using an E600 epifluorescence microscope equipped with a DXM1200 digital camera (Nikon, Mississauga, ON). Grayscale images were converted to pseudocolors using Photoshop software (version 23.1.1, Adobe, San Jose, CA).

The primary antibodies used in combination here have been tested separately resulting in similar labeling of the same neuronal structures. Removing the primary antibodies from the procedure while keeping everything else the same did not produce any labeling of neuronal structures in our material. The rabbit anti-DBH from this study was successfully used in zebrafish by other authors to label locus coeruleus neurons specifically^112,113^. In our material, only locus coeruleus neurons were labeled with this antibody.

Alternatively, the rabbit anti-DBH from this study was replaced with a rabbit anti-tyrosine hydroxylase (RRID: AB_390204, cat. number AB152, lot: 3031639, diluted 1:400, Millipore). The resulting labeling of the locus coeruleus neurons was similar in every aspect to the one obtained with the DBH antibody. The goat anti-ChAT used in this study has been used with success on a wide range of vertebrate species, including zebrafish, to label cholinergic neurons^49,113–115^.

#### Optical backfills using spatially-confined photoconversion of Kaede

Transgenic *Tg(vsx2:Kaede) z*ebrafish larvae expressing Kaede protein in V2a neurons were raised in darkness. At 4 dpf, larval zebrafish were anesthetized in 0.02% MS-222 and mounted dorsal side up in 1.5% in low melting point agarose. The Kaede protein in the rostral spinal cord (segments 4-9) was photoconverted by using a UV LED (405nm, Thorlabs, SOLIS-405C) for 2 minute-long illumination (500ms on and 500ms off pulsed illumination) through a 40X NA = 0.8 water immersion objective (power: 1.02 mW/mm2). This photoconversion procedure was repeated five times with intervals of 40 minutes. Digital mirror devices^116–118^ were used to confine the illumination in 2 dimensions from segment 4 to 9. Upon completion, the larvae were kept in the dark for 8 to 12 hrs to allow the photoconverted Kaede protein in the rostral spinal cord to travel back up to the hindbrain to label the soma of spinal projecting V2a neurons. The larvae were next imaged using a confocal microscope using a spinning disk head (CSU-W1, Yokogawa) on an upright microscope (Examiner Z1, Zeiss), using the 488 nm and 561 nm laser light sources (LaserStack, 3i Intelligent Imaging Innovations, Denver, USA). Images were acquired using a 20X N.A. = 1.0 DIC D=0.17 M27 75mm (Zeiss, no:421452-9880-000), using Slidebook software 6.0 (3i, Intelligent Imaging Innovations).

### Image analysis

All image analysis was performed using Fiji^111^.

### Calcium imaging experiments

For *in vivo* recording of calcium activity in V2a brainstem neurons (Fig. 4, 5), we screened the brightest fluorescent *Tg(vsx2:Gal4,UAS:GCaMP6s)* transgenic larvae at 3 dpf. On the day of the experiments, 6 dpf larvae were dorsally mounted as described above, with or without the tail left free to move according to the experiment’s needs. For experiments relying on MLR stimulations via single pulses, larvae were first paralysed by bath application of 1 mM α-Bungarotoxin solution (Tocris) diluted E3 medium for 3–6 min. Calcium imaging activity was recorded using a confocal spinning microscope (CSU-W1, Yokogawa) on an upright microscope (Examiner Z1, Zeiss; LaserStack, 3i Intelligent Imaging Innovations). Images were acquired using a 20X N.A. = 1.0 objective (Zeiss, no:421452-9880-000), using the Slidebook software 6.0 (3i, Intelligent Imaging Innovations). Images were acquired at either 16 Hz (Fig. 4) or 10 Hz (Fig. 5) with a field of view of 512 × 512 px and 0.666 μm/px pixel size. We choose to record planes representative of the different anatomical regions we imaged from, corresponding to planes at depth 182, 225, 257 and 288 μm in the *mapzebrain* brain atlas. After imaging calcium activity in each plane, we systematically recorded an anatomical z-stack of the entire hindbrain in order to localize the site of stimulation.

### Calcium imaging data analysis

#### Calcium imaging movie pre-processing

For all analysis of the fluorescence movies, we used the software *suite2p*^119^ (https://github.com/MouseLand/suite2p, v.0.8.0). The pipeline first corrected the movies for motion artifacts in 2D using rigid and nonrigid registration. Most regions of interest (ROIs) were automatically identified by the pipeline, and we manually added ROIs on cells that were not detected, including the soma of neurons that did and did not show activity during the recording. Then, using a custom-built Python script, for each ROI: *1)* we corrected the raw fluorescence trace by subtracting the neuropil signal (i.e., sum of surrounding pixels’ fluorescence, with a factor n = 0.7); *2)* we calculated the DF/F(t) = (F(t) - F0) / F0, where F(t) is the fluorescence at time t and F0 is the cell baseline fluorescence (median fluorescence value selected on a 3 s-long period of inactivity); 3) we estimated the noise, as the standard deviation of the signal during the same 3-s long period of inactivity, which was further used to set the recruitment threshold of the cell; 4) the DF/F was filtered using an average running filter with 3 time steps. For visualization purposes and regression analysis, a low pass filter was applied instead. In the tail-free experiments, when the motion artifact was larger and the 2D registration was not satisfactory, we removed the artifactual values by adding non existent values for frames in which the correlation to the mean image was low (we used 3 lowest percentile of correlation in the suite2p ops[‘corrXY’] output). In order to run correlation and regression analysis, we linearly interpolated the missing values to process continuous time series. Neurons were assigned to anatomical groups defined in Extended Data Fig. 2 based on their rostro-caudal position in the hindbrain and the manually-assigned boundaries between pontine, retropontine and medullary regions. For **Fig. 4**, each neuron was assigned to either ventral or dorsal group based on its normalized depth location in the *mapzebrain* atlas, using a dorso-ventral limit depth of 100 μm for pontine and retropontine regions and 80 μm for the medulla.

#### Registration of all calcium imaging planes across experiments

To compare recruitment of V2a neurons across fish, we registered ROIs from the calcium imaging data of all fish to a shared brain space (corresponding to one reference fish illustrated in **Fig.4** and **Fig.5**), as such: 1) anatomical boundaries, including the midline and the boundary between medulla and retropontine region-between r6 and r7, were visually identified (see **Extended Data Fig. 2**), and shifted to match the boundaries of the reference fish; 2) The same shift was then applied to each ROI’s position in order to extract a normalized X and Y position in the reference fish; 3) We estimated for each plane its normalized depth by manually comparing to which depth the plane corresponded the most to the *mapzebrain chx10:GAL4* fish space. The normalized coordinates were subsequently used for mapping neurons position in 3D.

#### Recruitment analysis in paralyzed transgenic larvae

In order to define whether a cell was recruited following an electrical stimulation in **Fig. 4**, we tested whether its calcium trace significantly increased above baseline. To do so, we checked whether the maximal DF/F value between the stimulation start and 3 seconds after was above an arbitrary threshold defined as the baseline plus 3 times the noise (standard deviation of the signal during the baseline). The baseline, for each cell at each stimulation, was defined as the mean signal over 1.5 seconds before the stimulation started. We defined ‘unresponsive cells’ as cells whose calcium trace did not reach threshold in the 3 s for any stimulation. Reliably-responding cells were the ones that reached the recruitment threshold systematically once a stimulation intensity was reached (**Fig. 4b**). Cells were classified as not reliably responding to the stimulation otherwise. For all cells that exhibited a calcium transient in response to the MLR stimulation, we computed the rising slope of the calcium trace as:

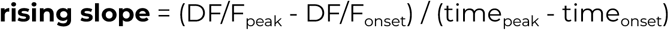

in which time_onset_ is the time when DF/F starts rising after the stimulation and time_peak_ is the time where the calcium transient peaks. As filtering calcium traces would induce a lag in the signal, we computed these time points on the raw DF/F. These time points were semi-automatically defined for all exhibited responses (see https://github.com/mathildelpx/Carbo-Tano_Lapoix_2022/notebooks/fig5_kinematicsAnalysisAuto.ipynb for detailed algorithm). The canonical V2a RSNs (RoM2, RoM3, MiV1, MiD2i, MiD3i) were named based on previous identifications^37,67^.

#### Recruitment analysis in tail-free larvae

We investigated which V2a neurons were specifically recruited during forward swimming episodes by comparing their activity during forward versus struggle episodes. To do so we extracted the maximum DF/F between 0.5 seconds before the start and 2 seconds after the end of the episode. As the max DF/F during a given episode could be influenced by the residual increase from a previous swimming episode, we corrected the max DF/F value by subtracting a normalized baseline (median DF/F of the 300 ms preceding the beginning of the episode). In these conditions, a cell was recruited if its corrected maximum DF/F reached 5 times the noise (defined as above in the preprocessing section).

#### Forward activity index and identification of a forward cluster based on calcium activity

In order to identify neurons recruited during forward episodes elicited by MLR stimulation, we clustered neurons using the calcium activity using an agglomerative clustering based on the python package *AgglomerativeClustering* from *scipy.cluster* library. We manually selected an arbitrary number of clusters (3 to 6) such that at least one cluster was specifically recruited during forward swimming only. To quantify that a group of neurons was indeed more active during forward swims, we computed for each neuron a *forward activity index* as:

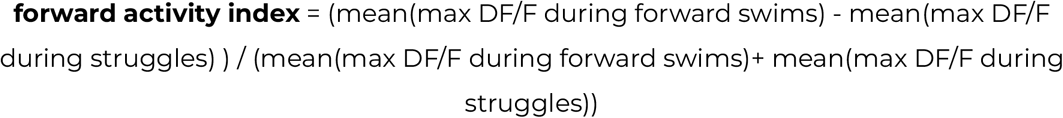

We verified for each trial that the cluster of neurons assigned as “forward” had the highest forward activity index.

#### Linear regression between max DF/F and number of oscillations

We used the package linregress from scipy.stats to model the max DF/F using the natural log of the number of oscillations. Here, we used a logarithmic transformation to get rid of the effects of the outliers in the distribution of the number of oscillations.

#### Building motor regressors

In order to compare neuron’s activity to motor activity, we built multiple motor regressors defined as fictive calcium traces modeling distinct kinematics parameters. Specifically, we first built four time series describing the tail beat amplitude, the tail beat frequency and for each of them, the binary value (0,1) based on their positive differential reflecting their increase over time. These were built as follow -

- <u>Instantaneous tail amplitude</u>: we resampled the tail angle trace (acquired at 300Hz) to the corresponding sampling rate of calcium imaging (10Hz), taking for each calcium imaging frame the maximal (in absolute) tail angle value.
- <u>Instantaneous tail beat frequency</u>: we calculated the mean tail beat frequency of all the cycles happening during each calcium imaging frame.
- <u>binary value of the positive derivative of the kinematic variables</u>: for each parameter computed above, we computed a binary variable equal to 1 when the parameter was increasing over time and equal to 0 the rest of the time. To do so, we used a Heaviside step function on the derivative of the kinematic parameter after filtering with a running average filter using 5 frames.

We then built the motor regressor concolving these time series with a GCaMP6s kernel modeled as negative exponential signal of decay 1.5 s.

#### Multiple linear regression to explain calcium trace with motor regressors

As each neuron could drive multiple kinematic parameters, we examined whether calcium activity could be explained by the four motor regressors described above, using multiple linear regression to model each neuron’s calcium trace DF/F, at each time step t, as:

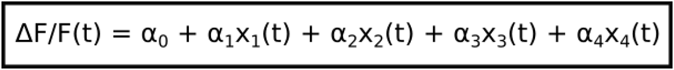

with x1 the regressor for the tail beat amplitude; x2 the regressor for the binary value of the increase in tail beat amplitude; x3 the regressor for the tail beat frequency; x4 the regressor for the binary value of the increase in tail beat frequency.

To fit the model on the data, we used the z-score of each kinematic parameter. We used a linear model optimized using ordinary least squares. The model aims to predict the calcium activity using a linear combination of the coupling coefficients (α1, α2, α3, α4) minimizing the error between the data and the model. We performed permutation tests to assess significance of a kinematic parameter in predicting the DF/F values (for significance, α needed to be positive and the p-value below 0.05). Overall, the kinematic parameters significantly explained respectively: iTBF, 59/89 neurons, iTA, 17/89 neurons; positive derivative iTBF, 6/89 neurons; positive derivative iTA, 32/89 neurons.

### Data visualization and statistical analysis

Analysis was done either in R^120^ (v4.0.4) using the tidyverse package^121^ (v1.3.0) or in Python 3.7. Initial figures were done either in R (v4.0.4) using the *tidyverse* package (v1.3.0) or using the *matplotlib^122^* (v3.3.2) and *seaborn*^123^ (v0.11.0) Python libraries. Figures were finally assembled in Inkscape (version 1.1, https://inkscape.org/). All statistical tests applied are specified in the legend of the corresponding figure. Statistical tests were performed in R (v4.04), except for the multiple linear model used in **Fig. 5**, for which we used the OLS function from the library *stats.models* developed for Python (https://github.com/statsmodels/statsmodels).

